# Movement during path integration discriminates age in a virtual-reality navigation task

**DOI:** 10.64898/2026.11.24.768162

**Authors:** Annie Kim, Gurleen Kaur, Afsoon G. Mombeini, Adam W. Lester, Manu S. Madhav

## Abstract

Healthy ageing brings about changes in our cognitive capabilities and how we interact with our environment. These changes are particularly evident in the way we approach complex tasks of daily living such as navigation. To understand how ageing affects different cognitive processes underlying navigation, we built a virtual reality task where 54 participants (ages 18-84) navigated a maze-like environment. They were then asked to point to their start locations and the locations of hidden distal landmarks. We computed error metrics based on the accuracy of their responses, and also tracked the continuous position and orientation of their head, hand and virtual body. Metrics indicative of path integration, i.e. keeping track of position by summing up motion signals, were affected by age. In particular, the angular error of pointing back to the start location shows a 1.06% increase per year of age. Older adults were not only less accurate at path integration but also, less consistent from trial to trial, which is what would be expected if ageing makes self-motion signals noisier. Critically, we found that classification of age group based on continuous movement kinematics was more accurate than classification based on pointing error metrics. A support vector machine (SVM) trained on 20 movement features discriminated age group with 80.7% balanced accuracy, which decreased to 73.4% when error metrics were added. Combining the SVM with a MiniRocket multivariate model trained on raw movement data further increased accuracy to 82.0%. Our results indicate that it is not how accurate you were, but rather how you move while you path integrate that is more discriminative of healthy ageing. It remains to be seen if continuous movement kinematics can also be used as a correlate of cognitive changes underlying neurodegenerative disorders like Alzheimer’s disease.

## 1. INTRODUCTION

### 1.1 Navigation Decline in Ageing and Neurodegeneration

Spatial navigation, our ability to represent, recall, plan and move through places in our environment, is a complex cognitive and sensorimotor ability. Navigation depends on the coordination of multiple sensory inputs, and their integration into multiple neural representations of space (Behrens et al., 2018; Cullen, 2019; Keshavarzi et al., 2023). These representations, termed a ‘cognitive map’, support relationships between ourselves and objects in our environment, and underlie spatial and episodic memory (Behrens et al., 2018; Epstein et al., 2017; Tolman, 1948). Egocentric representations encode locations relative to the observer (e.g. “the library is to my left”), whereas allocentric representations encode object-to-object relationships within a stable, world-centred frame independent of the observer’s position (e.g. “the café is north of the library”). Efficient navigation depends on transforming between these representations (Coughlan et al., 2018; Ekstrom et al., 2017; Ladyka-Wojcik & Barense, 2021). Path integration is the process in which self-motion cues – vestibular feedback, proprioception, and optic flow – continuously update position and heading (Savelli & Knierim, 2019; Segen et al., 2022). Since path integration accumulates errors in the absence of external input (Stangl et al., 2020), it must be periodically reset and calibrated against stable environmental features (Epstein et al., 2017; Jayakumar et al., 2019).

Ageing affects these different navigation-related computations selectively (Lester et al., 2017). Path integration decline is the most consistently reported deficit, particularly in ‘homing’ tasks where participants attempt to return to a start position (Adamo et al., 2012; Harris & Wolbers, 2012). Path integration deficits have been attributed primarily to noise in the velocity input, which accumulates with distance travelled rather than elapsed time (Stangl et al., 2020). These deficits have been associated with reduced entorhinal grid-cell-like representations (Moon et al., 2022; Stangl et al., 2018), which are able to predict path integration error in older adults. While allocentric memory is also less precise with age, strategy use and generalization across viewpoints can be preserved under immersive conditions (McAvan et al., 2021), and age differences are smaller for task paradigms that permit natural locomotion compared with desktop-controlled ones (Hill et al., 2024). In landmark processing tasks, which can involve both ego- and allocentric operations, older adults reorient less efficiently and show altered activity in scene-selective brain regions compared with younger adults (Bécu et al., 2020, 2023; Ramanoël et al., 2020).

This behavioural profile is of clinical interest because the circuits that subserve these navigation computations and decline earliest in ageing overlap with those affected earliest in Alzheimer’s disease (AD). Neurofibrillary pathology begins in the transentorhinal cortex before further spreading to limbic and neocortical regions (Braak & Braak, 1991), and entorhinal activity is disrupted even before neurodegeneration is detectable (Igarashi, 2023). These circuits also overlap with those for episodic memory, though as a behavioural marker, episodic memory discriminates incipient AD poorly, since it also declines in healthy ageing and is impaired in other dementias (Coughlan et al., 2018). Navigational measures have therefore been proposed as a more specific marker of early AD, one that also carries fewer cultural and educational biases. Path integration seems a particularly good candidate as it’s metrics can separate biomarker-positive from biomarker-negative mild cognitive impairment (MCI) more accurately than standard neuropsychological tests (Howett et al., 2019), predict AD risk in asymptomatic midlife adults more selectively than episodic memory or other spatial abilities (Newton et al., 2023, 2024), distinguish subjective cognitive decline from healthy controls (Segen et al., 2025), and predict subsequent cortical thinning in AD-vulnerable regions in cognitively unimpaired adults (Kawabata et al., 2026). While path integration deficits are also present in healthy ageing, they are more pronounced in early AD (Segen et al., 2022).

### 1.2 Virtual Reality as a tool for navigation assessment

Immersive virtual reality (VR) reconciles two goals that are difficult to satisfy together in real-world testing: experimental control and ecological validity. A head-mounted display (HMD) presents every participant with the same environment, landmarks, and path geometry, so that trials are reproducible across individuals and sessions, while still eliciting the visual self-motion cues that support natural navigation. Unlike desktop paradigms, where age differences may be confounded with unfamiliarity with keyboard or mouse control, some immersive setups permit physical rotation and even ambulation, adding vestibular and proprioceptive self-motion cues. Age differences in spatial memory are attenuated under these more naturalistic, cue-supported conditions (Hill et al., 2024; McAvan et al., 2021). VR systems can also record the continuous position and orientation of the head (HMD) and hands (controllers), yielding a stream of movement data that is much richer than discrete accuracy scores on which most navigation studies rely.

Navigation behaviour measured in virtual environments can distinguish healthy ageing from early neurodegeneration. Computer-based navigation performance tracks with preclinical AD as assessed by cerebrospinal-fluid biomarkers (Allison et al., 2016), and a VR test separated biomarker-positive MCI more accurately than standard neuropsychological measures (Howett et al., 2019). A recent systematic review of VR screening across task domains reported high sensitivity and specificity for MCI and early-stage AD, noting that the studies applying machine learning (ML) to behavioural output improved diagnostic accuracy, though the evidence base remains limited by small samples and methodological heterogeneity (Liu et al., 2026). A cognitive-motor account of navigational ageing argues that the movement produced during navigation is itself informative, reframing navigation as an ecologically valid cognitive-motor phenotype rather than a purely higher-order cognitive function (Hill & Ekstrom, 2025). Together these strands motivate an assessment that exploits the continuous movement captured by VR, rather than accuracy scores alone.

### 1.3 Gap: movement vs. metrics assessment

Most navigation assessments still reduce behaviour to a small set of hand-designed summary scores or metrics, such as path length, pointing error, and latency, much like explicit, structured cognitive testing. These metrics are easy to compute, but commit in advance to which aspects of behaviour are treated as informative. They may discard age- or disease-related structure that is present in the raw movement but not captured by the chosen summaries. Structured testing also adds participant burden and requires trained administration and scoring, which limits scalability. Although VR systems are capable of recording continuous movement data, only a minority of VR screening studies analyze the behavioural record, and those that apply ML to it report improved diagnostic accuracy (Liu et al., 2026).

An alternative is passive movement assessment: classifying participants directly from the continuous movement they produce while completing a task, without additional tests, targeted questions, or responses the participant knows are being scored. A movement classifier imposes no burden beyond the task itself, and it can in principle be embedded in VR applications that already log head and controller motion, whether for rehabilitation, training, or entertainment. In this study, we explore whether raw movement carries age-related information that hand-designed summaries miss, and whether a movement-based classifier and a hybrid movement-metrics classifier can recover this information.

## 2. METHODS

All procedures, materials and protocols were approved by the University of British Columbia Behavioural Research Ethics Board (BREB; Protocol #H21-02923).

### 2.1 Participants

Sixty-five community-dwelling adults and undergraduate students were recruited through advertisements on our laboratory website, departmental and community mailing lists, the REACH BC public health research platform, and the University of British Columbia Psychology Department Human Subject Pool (HSP). Community volunteers received $10 CAD as reimbursement of travel and parking costs, while HSP participants received two course credits. Eligible participants were 18 years or older, fluent in English, and had intact or corrected-to-normal vision and hearing. Individuals were ineligible if they reported a diagnosis of epilepsy or cognitive impairment, physical limitations affecting neck or arm movement, or a history of motion sickness or a diagnosed vestibular disorder during screening. All participants provided written informed consent and were also informed of their right to withdraw at any time.

Three rule-based exclusion criteria were applied after data collection. Participants were excluded if they (a) completed fewer than 15/30 experimental trials, leaving insufficient data to estimate within-participant trends; (b) had missing Corsi span or Penn Matrix Analysis Test (PMAT) scores due to online testing platform failures; or (c) scored below the standard clinical threshold on the Montreal Cognitive Assessment (MoCA < 26). These exclusion criteria were not mutually exclusive: six participants completed fewer than 15 trials, three had missing cognitive data, and four scored below the MoCA threshold, yielding 10 unique rule-based exclusions. Of the six participants completing fewer than 15 trials, four withdrew because of persistent cybersickness, one because of fatigue, and one because of a headset malfunction. One additional participant was excluded post-hoc, from session notes indicating that this participant had substantial difficulty understanding the task instructions and required supplementary explanation from the researcher partway through the session. In total, 11/65 participants were excluded from analysis.

The final analytic sample thus consisted of 54 adults (30 female, 24 male) aged 18 to 84 years (M = 43.30, SD = 22.63) contributing a total of 1,633 trials. (1,568 in blocks 1-6) Since middle-aged adults were underrepresented, the age distribution was approximately bimodal (Supp. Fig. 1) and therefore participants were classified as younger (< 55 years; n = 33, M = 27.21, SD = 11.59) or older (≥ 55 years; n = 21, M = 68.57, SD = 6.64) for robustness analyses. The 55-year threshold was selected after inspecting the observed age distribution and was not pre-specified. No participant in the analytic sample was aged 55 to 58, so any threshold within this gap yields identical groups. Sample characteristics are detailed in Table 1.

**Table 1.** Sample Characteristics by Age Group.

| Characteristic | Overall | Younger | Older | Test | p |
|---|---|---|---|---|---|
| N | 54 | 33 | 21 | — | — |
| Age (years) | 43.30 (22.63) | 27.21 (11.59) | 68.57 (6.64) | t(51.6) = 16.65 | < .001 |
| Trials contributed | 29.04 (3.19) | 29.58 (2.44) | 28.19 (4.03) | t(29.4) = -1.42 | .167 |
| Education (years) | 17.41 (3.60) | 16.27 (3.56) | 19.19 (2.93) | t(48.5) = 3.28 | .002 |
| Corsi span | 5.63 (1.44) | 6.09 (1.42) | 4.90 (1.18) | t(48.3) = -3.32 | .002 |
| PMAT (/24) | 17.74 (3.93) | 18.91 (3.08) | 15.90 (4.47) | t(32.0) = -2.70 | .011 |
| Gaming experience (0-4) | 1.37 (1.31) | 1.79 (1.34) | 0.71 (0.96) | t(51.2) = -3.43 | .001 |
| VR experience (0-4) | 0.74 (0.62) | 0.97 (0.59) | 0.38 (0.50) | t(47.6) = -3.95 | < .001 |
| State anxiety (STAI-S) | 5.98 (1.69) | 6.33 (1.99) | 5.43 (0.81) | t(45.8) = -2.32 | .025 |
| Trait anxiety (STAI-T) | 8.56 (3.27) | 9.45 (3.33) | 7.14 (2.67) | t(49.1) = -2.81 | .007 |
| Depression (CESD-10) | 6.70 (5.32) | 8.82 (5.40) | 3.38 (3.06) | t(51.5) = -4.72 | < .001 |
| MoCA (/30) | — | not administered | 28.76 (1.34) | — | — |
| Sex, n female (%) | 30 (55.6%) | 19 (57.6%) | 11 (52.4%) | Fisher exact | .783 |
| Family history of cognitive impairment, n (%) | 22 (40.7%) | 10 (30.3%) | 12 (57.1%) | Fisher exact | .087 |
| Self-reported sun use, n (%) | 15 (27.8%) | 7 (21.2%) | 8 (38.1%) | Fisher exact | .220 |
| Ethnicity, n (%) |  |  |  | — | — |
| Asian | 19 (35.2%) | 17 (51.5%) | 2 (9.5%) |  |  |
| Middle Eastern | 1 (1.9%) | 0 (0.0%) | 1 (4.8%) |  |  |
| Mixed | 5 (9.3%) | 5 (15.2%) | 0 (0.0%) |  |  |
| South Asian | 2 (3.7%) | 2 (6.1%) | 0 (0.0%) |  |  |
| White | 27 (50.0%) | 9 (27.3%) | 18 (85.7%) |  |  |

Notes: N = 54. Continuous variables are M (SD). Group comparisons use Welch's t-test (continuous) or Fisher's exact test (categorical) and are descriptive: they are reported without multiplicity correction and were not used to select covariates. MoCA was administered only to participants aged 59 years and older; the older sample is therefore cognitively screened (MoCA >= 26). Ethnicity is reported descriptively with no group comparison, as it is not used in any analysis.

Prior familiarity with virtual-reality and gaming interfaces is a plausible confound in age-group separation: older and younger adults may differ in interface fluency as well as in spatial navigation. Both measures were recorded at intake as two self-report items (VR experience and gaming experience). Both differ by age group (Table 1). Neither was included as a feature in any classifier, and no analysis tested whether classifier performance is attributable to interface familiarity rather than to navigation. This potential confound is addressed in Section 4.8.

### 2.2 Demographics and Cognitive Measures

Demographic and mood questionnaires were administered via the University of British Columbia Survey Tool (Qualtrics, LLC. Provo, UT U.S.A.) and distributed the day before the experimental session to reduce in-lab testing time. Cognitive assessments were administered in-person on the day of the session by trained research personnel.

#### Demographics Questionnaire

A custom questionnaire recorded biological sex, age, ethnicity, years of education, medical history (including family history of cognitive impairment), and self-reported rate of prior experience with virtual reality and video gaming, each on a five-point scale from none at all (0) to a great deal (4).

#### Depressive Symptoms

The 10-item Center for Epidemiologic Studies Depression Scale (CES-D-10; (Andresen et al., 1994)) assessed depression symptom frequency over the last week on a 0–3 scale. Two positively worded items were reverse-coded, and the items were summed (range 0–30), with higher scores indicating greater symptom severity.

#### Anxiety

Abbreviated five-item State (STAI-S-5) and Trait (STAI-T-5) subscales of the State-Trait Anxiety Inventory (Zsido et al., 2020) were administered. Items were rated from not at all (1) to very much so (4), yielding subscale scores ranging from 5 to 20.

#### Global Cognitive Screening

The Montreal Cognitive Assessment (MoCA; (Nasreddine et al., 2005)) was administered to participants aged 59 years and older by researchers who had completed the official MoCA training certification. Because no participant in the analytic sample was aged 55 to 58, the cognitively screened participants coincide with the older group as defined for robustness analyses. Scores were used solely for screening and were not entered into any statistical model. All 21 older adults in the analytic sample completed the MoCA.

#### Non-verbal Reasoning

The Penn Matrix Analysis Test (PMAT; (Gur et al., 2010)) was administered through the Penn Computerized Neurocognitive Battery (PennCNB; (Moore et al., 2015)). Participants viewed a patterned matrix and selected the best-fitting cell from five different options, across 24 trials of increasing difficulty. The test was automatically terminated after five consecutive errors. Performance was scored as the number of correct responses. PMAT was retained as a sample descriptor and was not entered as a covariate.

#### Visuospatial Working Memory

A computerized Corsi block-tapping task (Kessels et al., 2000) administered via PsyToolkit (Stoet, 2010, 2017), presented nine blocks that illuminated in sequence, beginning at a length of two, and participants were required to replicate the sequence. The sequence length increased after each correct trial and was repeated after each error; the task terminated after two consecutive errors at the same length. Corsi span was defined as the longest sequence correctly reproduced.

#### Exit Survey

After the VR task, participants rated the clarity of instructions and controls, reported any cybersickness symptoms, and described any strategies used, including reliance on environmental cues. Participants who explicitly mentioned using the sun as a directional cue were classified as sun users (n = 15) and this classification was used as an exploratory covariate. Since this relied on unprompted mentions in free-text responses rather than a targeted question, it is a conservative indicator of sun use.

### 2.3 VR spatial navigation task

#### VR Hardware & setup

The task was administered using the Quest 3 HMD (Meta, CA, U.S.A.; 120 Hz refresh rate; 2064 × 2208 pixels per eye), connected wirelessly to the laboratory computer via the Meta AirLink protocol. During the task, participants were seated in a swivel chair with wheel stoppers positioned centrally in the testing room, and rotated their bodies freely to change heading, engaging vestibular and proprioceptive cues, and visual cues through the headset (Fig. 1a). Translation was controlled by the thumbstick on the controller, while landmark selection was performed by aiming the right controller and pressing the trigger. Task instructions were displayed on a virtual panel attached to the left controller; participants could raise their left arm into their field of view to review instructions. Participants were instructed to use their right controller for pointing tasks and use the left controller only for referencing instructions.

**Figure 1.**
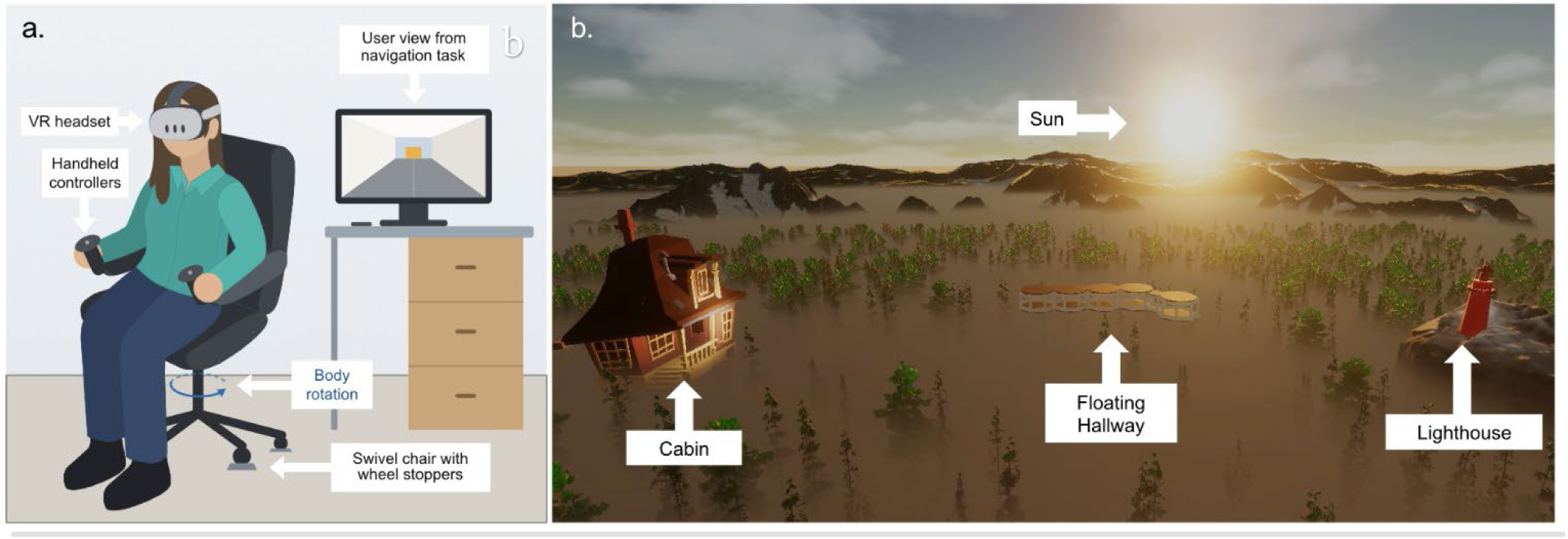
(a) Experiment setup. A wireless VR headset provided visual feedback, while handheld controllers enabled translational movement. Participants physically rotated their bodies to change heading direction. (b) Virtual environment. Two salient landmarks, a lighthouse and cabin, were in fixed locations, along with the sun as a directional reference. The participants navigated in a floating hallway, whose shape changed in every trial.

#### Virtual Environment

The virtual environment (Fig. 1b) was built in the Unity Game Engine (Unity Technologies, CA, U.S.A., v.2021.3.17f1) and consisted of naturalistic terrain and lighting, ambient sound, and the sun as an optional global directional cue. These environmental features, along with two key landmarks – a lighthouse and a cabin – remained fixed in position throughout the task, serving as stable allocentric reference points. The navigation task took place within a floating hallway assembled from interconnected octagonal cells, whose configuration and position varied across trials. The layout of these cells was designed to mimic our rodent Omniroute maze (Lester et al., 2025), for future cross-species behavioural comparisons.

#### Task Design

Prior to our ‘Memory Trails’ task, participants received verbal instruction, an interactive video tutorial, and three practice trials in a fully visible straight hallway, followed by a narrated demonstration of a complete trial and one interactive practice trial.

Each trial consisted of six sequential stages within a single continuous episode (Fig. 2a):

1. **Environment encoding:** From a starting room with panoramic windows, participants viewed and memorized the locations of the lighthouse and cabin. The sun was also visible but not explicitly stated to participants as informative.
2. **Path integration**: Participants exited the room into a fog-obscured hallway that occluded all external landmarks and traversed a predefined path to a cone marking the endpoint. Heading changes relied on self-motion cues from physical body rotation in the swivel chair.
3. **Start pointing**: At the end of the path with the surrounding walls removed and no landmarks visible, participants indicated the remembered start location by placing a blue pin on the cell corresponding to their perceived start position, using a to-scale octagonal grid overlaid on the environment.
4. **Reorientation**: The true start location (marked with a green holographic pin) and the sun were revealed, and participants acknowledged the true start. This stage provided external cues to correct accumulated path integration errors.
5. **Landmark pointing**: With both reorientation cues visible, participants indicated the direction of the lighthouse then the cabin, by placing holographic markers. The placed lighthouse marker remained visible during cabin placement, and the order was fixed across trials.
6. **Feedback**: The full environment was revealed, and participants received an accuracy score. Participants acknowledged the true landmark positions prior to beginning the next trial.

**Figure 2.**
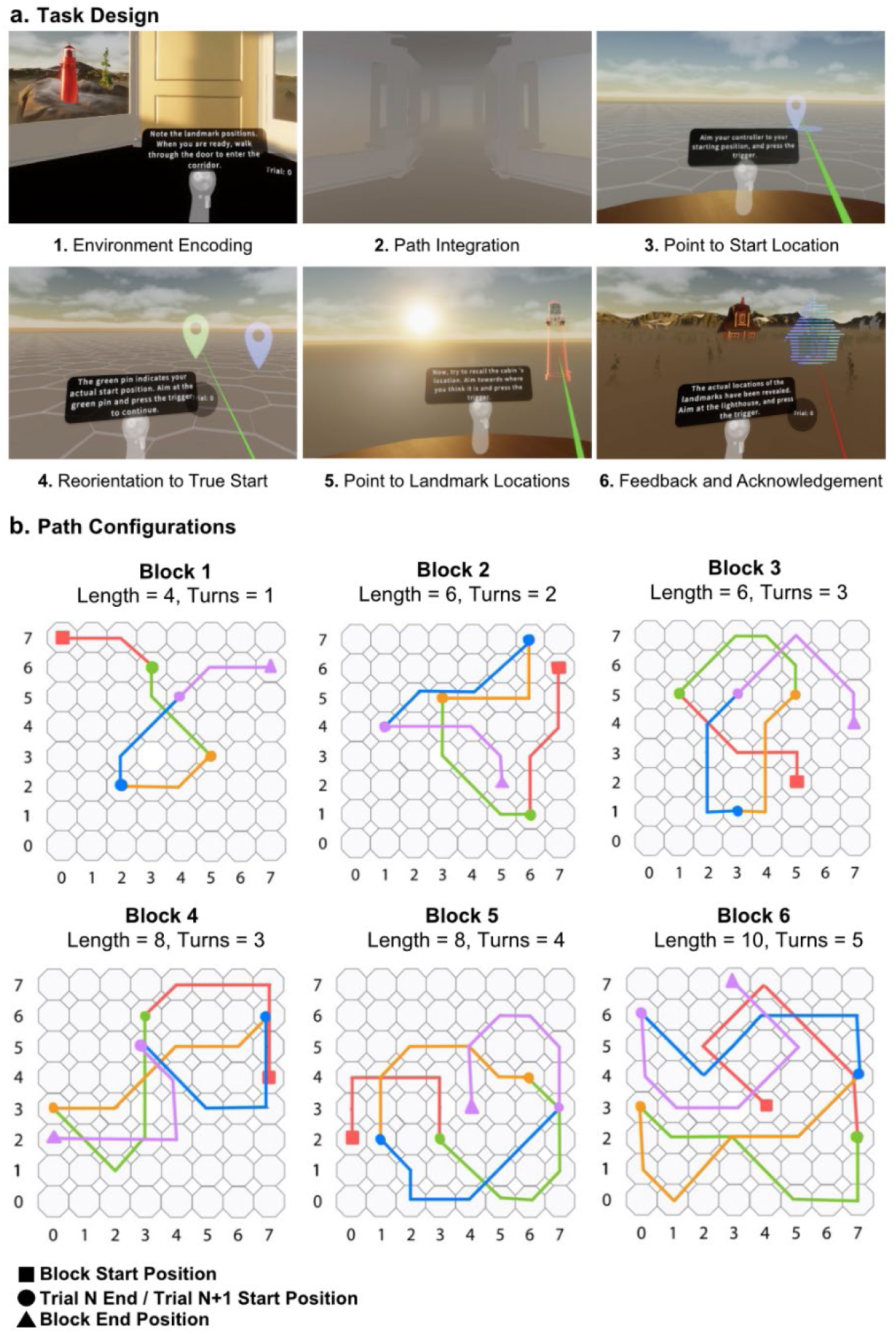
(a) ‘Memory Trails’ task design. Each trial consisted of six distinct stages (1-6). Participants (1) pointed to the location of the two landmarks, (2) walked through a hallway while surrounded by fog, (3) pointed back to their start location, (4) was shown and acknowledged the true start location, (5) pointed to the two hidden landmarks, and (6) were shown the true location of the landmarks. (b) Path configurations. The maze was arranged in a 8×8 grid of octagonal rooms. 30 trials (colored lines) were grouped in 6 blocks of increasing complexity (length, number of turns). The end position of each trial was the start position of the next trial. The order of trials, blocks and the trajectories themselves were all consistent across participants.

The task comprised 30 trials divided into six blocks of five. Trials were chained such that the end position of trial n became the start position of trial n + 1, with no teleportation or reset. Across blocks, the path length and number of turns increased systematically; all trials within a block matched in length and turn count but differed in spatial configuration (Fig. 2b). Turns were quantified in 45° increments, such that one 90° turn is counted as two turns; no paths included turns larger than 90°. Since block order was fixed and path complexity increased monotonically, complexity and time-on-task are colinear in this design and cannot be separated.

A seventh block, repeating the path configurations of block 1, was introduced partway through data collection, and was offered to all subsequent participants. Thirteen of the 54 participants in the analysis sample (6 younger, 7 older) completed both blocks 1 and 7, and these trials were compared within participants as an exploratory analysis.

### 2.4 Data acquisition and preprocessing

#### 2.4.1 Event Data

Discrete event data including stage transition timestamps, fixed and participant-selected landmark coordinates, and trial path configurations, were logged as one JavaScript Object Notation (JSON) file per participant.

For each trial, the true and participant-indicated bearings of the start, lighthouse, and cabin were computed from the participant’s endpoint position in world coordinates, so that all angles are independent of the participant’s own heading. Trials were screened for accidental mis-clicks (i.e., double-clicking of the trigger that placed the next hologram marker on top of the one already visible) based on two rules – a pointed lighthouse coinciding with the visible true start marker, and a pointed cabin coinciding with the visible pointed lighthouse, using an angular threshold of 0.25°. A similar artefact cannot arise during the start-pointing, as no holographic marker is present on the grid at the time of the start response phase. An additional 15 trials were flagged as mis-clicks through manual review of session notes. Flagged trials contributed no data to any outcome (34 of 1,633 trials, 2.08%), so that trial counts are identical across outcome measures and resulted in a total of 1,599 usable trials across all seven blocks. Of these, 1,535 usable trials fell within the six analysis blocks and entered the statistical models.

#### 2.4.2 Continuous Data

Continuous position and orientation were logged from the HMD and both hand controllers at a nominal rate of 66.7 Hz (median inter-frame interval 15 ms), with timestamps recorded to millisecond resolution. However, the sampling was irregular: the mean inter-frame interval was 20.0 ms (interquartile range 14 to 28 ms), reflecting intermittently dropped frames, and the effective mean sampling rate across the dataset was ∼50 Hz. Each frame recorded a timestamp followed by 3D position (three values) and quaternion orientation (four values) for each of the three tracked rigid bodies (HMD, right controller, left controller), giving 21 kinematic values per frame. Positional values are expressed in Unity world units, which correspond to metres in the virtual environment. Duplicate consecutive frames, arising from logging overlap and identifiable by identical timestamps and values, were removed. All rate-based quantities were subsequently computed from true per-frame time intervals rather than assuming uniform frame spacing.

ML analyses (Sec 2.6) used kinematic features from the HMD and the right controller (14 channels) only. The left controller was excluded because participants used it only to raise the instruction panel into view, so its movement reflects instruction-checking rather than navigation. Additionally, data was restricted to the first 16 trials per participant, since 16 was the largest trial count reached by every participant in the sample. Since session length was shorter on average among older adults, an uncapped analysis would have weighted younger participants more heavily.

Two intervals were defined in each trial: the *follow phase* and the *point phase*. The follow phase corresponds to stage 2, beginning when the participant exits the encoding room into the fog-obscured hallway and ending when they reach the cone marking the endpoint of the path. It is the only interval in which the participant translates through the environment, and the only one in which position must be maintained from self-motion cues alone. The point phase begins at that same cone and ends when the cabin marker is placed, spanning stages 3 through 5: the start-pointing response, the reorientation in which the true start and the sun are revealed, and the lighthouse and cabin responses. During the point phase, the participant remains at the path endpoint throughout, so the movement recorded is head and controller motion rather than translation. The two intervals therefore differ both in what the participant is doing and in what the recorded movement can reflect, which is why kinematic features are computed separately for each rather than pooled across the trial.

### 2.5 Error metrics modeling

The study was designed around error metrics computed from Event data (Sec. 2.4.1) and modeled using statistics across age groups and blocks. Analysis of error metrics as a continuous function of age was performed using Generalized Linear Mixed Models (GLMMs).

#### 2.5.1 Error metrics

Five angular error metrics were derived per trial, each computed in both signed (directional) form and absolute (magnitude) form (Fig. 3). Absolute errors served as the outcome for all models, and signed errors were used for circular descriptive statistics and for the exploratory error coupling analysis.

**Figure 3.**
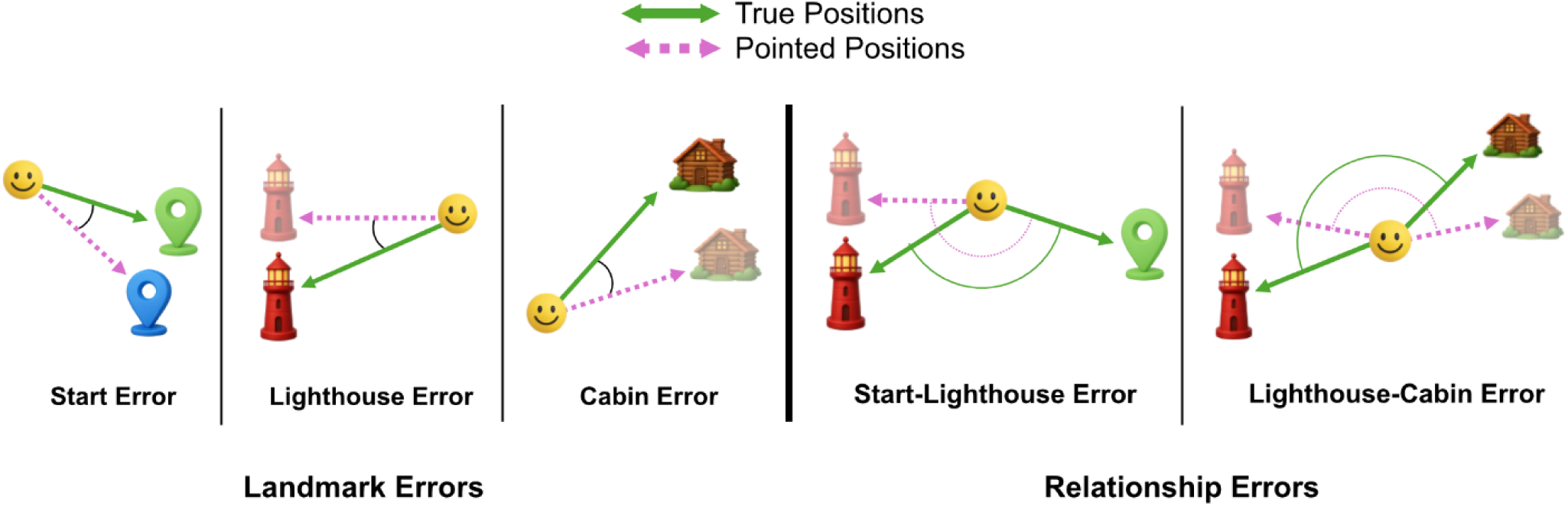
Error metrics. (left) Three Landmark errors measured the errors between pointed (lighter) and true (darker) locations of the start, lighthouse and cabin, respectively. (right) Two Relationship errors measured errors between the relative angles between the pointed and true landmarks.

**Landmark Errors** quantify the angular discrepancy between the true and pointed landmark directions. All directions (θ) were computed from the participant’s end position and measured counterclockwise when viewed from above. All angular arithmetic was performed circularly, with landmark errors wrapped to [-180°, 180°). Signed errors were computed such that positive values indicate pointing too far right, and negative values indicate pointing too far left, of the true direction.

1. **Start Error** is the discrepancy in the remembered start location, indexing egocentric path integration from self-motion cues.

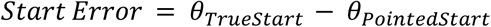
2. **Lighthouse (LH) Error** is the discrepancy in the lighthouse following reorientation, indexing cue-based reorientation.

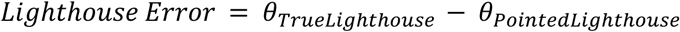
3. **Cabin Error** is the discrepancy in the cabin, indexing allocentric scene memory.

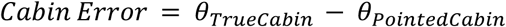

**Relationship Errors** quantify the fidelity of the angular separation between a pair of landmarks, independent of their absolute global direction, with positive values indicating overestimation and negative values indicating underestimation of the separation. Each separation |*θ_a_* − *θ_b_*| denotes the smallest angular separation, in [0°, 180°]:

1. **Start-LH Error** compares the angular separation between the true start and the pointed lighthouse against the true separation.

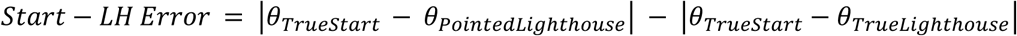
2. **LH-Cabin Error** compares the angular separation between the pointed lighthouse and pointed cabin against the true separation.

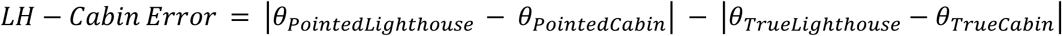

Since the true start marker was visible during lighthouse placement, and the pointed lighthouse marker was visible during cabin placement, each relationship error is anchored to the reference the participant could actually see. As relationship errors are computed from unsigned angular separations, they index the accuracy of the perceived gap between two landmarks but not which side of the reference landmark it was placed (e.g., with a true Start-LH separation of 60°, a lighthouse placed 90° to the left or 90° to the right of the true start yields the same Start-LH Error of +30°).

Start-LH Error and LH Error are closely related by construction. Both are computed from the pointed lighthouse direction, but Start-LH Error measures it against the true start rather than the true lighthouse. In most trials, pointing a given number of degrees away from the true lighthouse widens or narrows the Start-LH separation by the same number of degrees, so the two absolute errors are identical. However, the absolute values of the Start-LH Error and LH Error differ when the pointed lighthouse direction crosses the true start direction or its 180° opposite. Because an angular separation is bounded between 0° and 180°, the separation folds back instead of changing by the full pointing error. Accordingly, participant-level absolute errors were near redundant in the present sample (Spearman *ρ* = .97; Table S11a). In their signed forms, however, the two metrics describe different properties of the same response: signed LH Error indicates the direction of the pointing error (left or right of the true lighthouse) whereas signed Start-LH Error indicates whether the Start-LH separation was over-or under-estimated. For example, pointing too far right of the true lighthouse widens the Start-LH separation when the lighthouse lies to the right of the start, but narrows it when the lighthouse lies to the left. The same pointing error can therefore produce a positive or a negative Start-LH Error, depending on where the lighthouse was placed relative to the start on that trial. Consistent with this, the signed metrics were essentially uncorrelated (circular *r* = −.02; Table S11b). Start-LH Error is retained for completeness but should not be interpreted as an independent construct.

Absolute Lighthouse and Cabin Errors were also closely correlated (*ρ* = .93; Table S11a), although, unlike Start-LH and LH Error, they share no common terms by construction. In their absolute form, the correlation matrix (Table S11a) suggests that the five error metrics form approximately three clusters: Start Error; Lighthouse, Cabin, and Start-LH Errors; and LH-Cabin Error.

#### 2.5.2 Error metrics analyses

Analyses were conducted in Python 3.13.5 and R 4.6.1, with R called from Python through *rpy2 v3.6.7*.

##### Descriptive Analyses

Participant-level means of each absolute error metric were computed across blocks 1-6, and age-group differences were summarised with Hedges’ g. These descriptive comparisons are reported without multiplicity correction and are intended to characterize the sample rather than to test hypotheses. Block-wise descriptives were computed by first averaging within each participant and block before aggregating.

##### Trial-to-Trial Variability Analysis

To investigate trial-to-trial response variability, participant-level standard deviations of each absolute error metric were computed across blocks 1-6. Age group differences in the mean standard deviations were tested with two-sided Monte Carlo permutation tests (100,000 resamples), randomly re-assigning each participant to age groups and recomputing the group difference on each permutation. Percentile bootstrap confidence intervals (10,000 resamples) were computed for the group difference in degrees, and effect sizes are reported as Hedges’ g, with positive values indicating greater variability in older adults. Holm correction was applied across the five error metrics. The association between the number of trials a participant completed and their standard deviation was also examined using Spearman correlation, to test whether unequal session length influenced the variability of estimates.

##### GLMM Model

Each absolute error metric was analyzed with a Generalized linear mixed model (GLMM) fitted by maximum likelihood in glmmTMB v1.1.14 (Brooks et al., 2017), using a Gamma distribution with a log link. This family was selected because angular error magnitudes are strictly positive, right-skewed (skew 0.72-1.42), and contained no exact zeros. Gamma was preferred to a lognormal distribution by the Akaike Information Criterion (AIC; (Akaike, 1974)) for all five metrics (Table S2a). A by-participant random intercept accounted for the non-independence of trials within participants.

The Gamma family ties residual spread to the fitted mean, so with constant dispersion, any age difference in mean error produces a proportionally equal age difference in residual spread. Therefore, each model also included a dispersion sub-model (primary: *formula = ∼Age_c*; binary: *formula = ∼ AgeGroup*) allowing the Gamma dispersion parameter to vary with age. Inclusion was assessed through likelihood ratio tests against a constant-dispersion specification (*formula = ∼1*), The dispersion component is estimated on a log scale, and its coefficients describe the Gamma dispersion parameter rather than variability in degrees. Model selection statistics and dispersion coefficients are reported in Table S2d. The primary model was defined as:

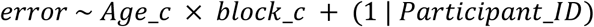

where *Age_c* is age in years, mean-centred within the sample, and *block_c* is block number minus 3.5 (block_c = block – 3.5), so that blocks 1-6 are coded as –2.5 to +2.5 and block_c = 0 falls at the midpoint of the session. This centering allows the age coefficient to describe the age effect at mid-session rather than at an extrapolated, non-existent block zero. Because the link is logarithmic, exponentiated coefficients are multiplicative ratios of the expected error. To interpret the per-year age coefficient alongside the group-based robustness analysis, age effects are additionally reported scaled across the observed 41.36-year gap between the mean ages of the younger and older groups.

##### Age Specification Robustness

Since the sample age distribution was approximately bimodal (Supp. Fig. 1), all primary models were refitted replacing continuous age with the binary younger/older classification, retaining an equivalent dispersion sub-model.

##### Estimated Marginal Means

Model-implied error magnitudes were obtained using *emmeans* v2.0.4 (Searle et al., 1980) on the response scale, evaluated at mid-session and, for the continuous-age models, at ages 27 and 69 (the group means, rounded). Marginal means were evaluated at a fixed block rather than averaged across blocks, because in the presence of age x block interactions, effects averaged across blocks would be difficult to interpret.

##### Multiplicity Correction

Holm correction was applied within families of five (one test per error metric) separately for the age effect, the block effect, the age × block interaction, the age-group difference in trial-to-trial variability, and the Watson U^2^ group comparison. Benjamini-Hochberg false discovery rate (FDR) control was applied across the 15 covariate tests in the adjusted models, across the pooled 10 tests for gaming and VR experience, and across the 25 tests in the exploratory covariate analysis. Family boundaries were determined by the authors after data collection and were not pre-specified.

##### Model Diagnostics

Model fit was assessed with simulation-based scaled residuals computed from 1000 simulations per model using DHARMa v0.5.0 (Hartig, 2026), comprising tests of uniformity, dispersion, outliers, and quantile deviation. Collinearity among adjusted-model covariates was assessed by variance inflation factors.

##### Sensitivity Analyses

Four sets of sensitivity analyses were conducted. Five participants (one younger and four older adults) withdrew before completing all six blocks. As this uneven attrition could bias age comparisons, the trial-to-trial response variability analysis was repeated on the completers-only panel of 49 participants (32 younger, 17 older). The GLMM models were also refitted on the completers-only panel. Attrition was examined by comparing the blocks 1-3 performance of completers and non-completers, and by comparing block-wise means and group effect sizes between the full sample and the balanced panel. In addition, models were refitted with a theoretically motivated covariate set (sex, education, and Corsi span). Finally, gaming and VR experience were each added to the adjusted model separately. Since education, Corsi span, gaming, and VR experience are themselves correlated with age, these models are reported as sensitivity analyses rather than as the primary estimate.

#### 2.5.3 Exploratory Analyses

1. **Group Comparison of Directional Distributions:** Since the GLMMs model the size of each error but discard its direction, signed errors were analyzed. For each participant, a mean direction was computed across their trials for each signed error, and these means were summarized by age group using circular statistics (mean direction, mean resultant length R, and 95% CI). R indexes the consistency of directional bias across participants rather than the precision of individual pointing responses. Age-group differences in signed error were tested with Watson’s U^2^ two-sample test, a nonparametric omnibus test of whether two circular samples are drawn from the same distribution - it does not assume unimodality or equal concentrations across groups. P-values were obtained by permutation, randomly re-assigning participants to age groups 10,000 times and recomputing U^2^ on each permutation. Holm correction was applied across the five error metrics.
2. **Practice Effects**: Since path complexity increased across blocks, the effects of practice and path configurations cannot be separated. A seventh block that repeated path configurations of block 1 was used to examine practice with path configuration held constant. For the 13 participants who had block 7, within-participant change from block 1 was tested with exact sign-flip permutation tests (all 8192 possible sign assignments) and percentile bootstrap confidence intervals (10,000 resamples). P-values were Holm corrected across the five metrics. Descriptive checks were also done to determine whether block 7 recipients differed from other participants in blocks 1-6 performance.
3. **Error Coupling:** Since the start, lighthouse, and cabin pointing responses in each trial were made within a single continuous episode, we examined whether errors were coupled across responses, which would indicate that the responses share a common rotational reference frame. For each participant, correlations were computed across their trials for the start–lighthouse, start–cabin, and lighthouse–cabin pairs: circular-circular correlation on signed errors for directional coupling, and Spearman correlation on absolute errors for magnitude coupling. Each participant’s correlation was treated as a single observation and tested against zero with sign-flip permutation tests (100,000 resamples), and between groups with two-sample permutation tests (100,000 resamples), with percentile bootstrap confidence intervals (10,000 resamples) and Holm-corrected p-values. The same tests were repeated comparing sun-users to non-sun-users, and within the sun-user subgroup alone. Since the sun becomes visible after pointing to the start and before landmark pointing, it offers a reorientation opportunity that could weaken coupling between the start response and the landmark responses.
4. **Covariates:** Sex (male as reference), z-scored years of education, and Corsi span constituted a theoretically motivated set of covariates, entered as main effects in adjusted models. Z-scored gaming experience and z-scored VR experience were each entered separately to the adjusted model as sensitivity covariates, as interface familiarity is age-correlated (Spearman ρ = −.50 and −.48 with age, respectively) and establishes a possible alternative account of age differences in the present VR-based task. Sun use, z-scored STAI scores, z-scored CESD scores, and family history of cognitive impairment (immediate family or close relative versus distant relative or none) were examined as exploratory covariates. Ethnicity and PMAT scores are reported descriptively in Table 1 and were not modelled.

### 2.6 Machine learning

Event and continuous data (Secs. 2.4.1 and 2.4.2) were used to train three machine learning (ML) approaches to age-group classification: a support vector machine (SVM), MiniRocket, and a late fusion of the two. For ML modeling, we used Python 3.10.20 with scikit-learn 1.7.2 (Pedregosa et al., 2011), sktime 1.0.0 (Löning et al., 2019), NumPy 2.2.6 and SciPy 1.15.3. We also used a constant random seed so that the analyses are precisely reproducible.

All models were evaluated by leave-one-participant-out cross-validation: each model was trained on 53 participants and tested on the one held out, repeated so that every participant served as the test case exactly once. Z-scoring of features, the MiniRocket transform, hyperparameter selection and the decision threshold were each fitted on the training participants of each fold separately, so no statistic used to classify a participant was derived from data that included that participant.

Older adults were treated as the positive class. Sensitivity is thus the proportion of older adults classified correctly and specificity the proportion of younger adults classified correctly. Balanced accuracy (BA) is the mean of the two, which corrects for the 33 to 21 group imbalance and has a chance value of 0.5 regardless of class proportions (Brodersen et al., 2010). The area under the receiver operating characteristic curve (AUC) was computed once per model from the held-out predicted probabilities of all 54 participants pooled across folds. AUC summarises how well a model ranks older above younger participants across thresholds, so it does not depend on a particular decision threshold (Bujang, 2026).

All models were assigned class labels by the same rule, so that their BAs are directly comparable. Each model produced a predicted probability of older-group membership for every held-out participant: for the SVM and MiniRocket this was the Platt-calibrated output (Platt, 2000), and for fusion it was the equal-weight average of the two models’ probabilities. Within each fold, a decision threshold was chosen on the training participants by maximising the Youden index, J = sensitivity + specificity – 1 (Youden, 1950), over the probabilities of the other 53 participants, and then applied to the held-out participant.

#### 2.6.1 SVM Feature extraction

Twenty-nine kinematic and task-performance features were extracted from each trial, computed separately for the follow phase (maze navigation) and the point phase (landmark identification) where applicable. One of these, the interval between the start-location pointing response and the participant touching the cone, was excluded from modelling: the event log records both at the same instant, so the interval is identically zero in 857 of 864 trials and the participant-level mean is undefined for 47 of 54 participants. The remaining 28 features can be organized into six groups based on the behavioural dimension they captured:

##### Rotational dynamics (2 values x 2 phases; 4 features)

*Mean and maximum angular velocity of head rotation* (degrees/s) were computed for both phases.

##### Heading behaviour (2 x 2; 4 features)

*Heading drift*, the standard deviation of the yaw of the forward heading vector (degrees), and *heading range*, the difference between its maximum and minimum (degrees), were computed for both phases.

##### Movement smoothness and pausing (2 x 2; 4 features)

*Smoothness* was quantified as the absolute angular acceleration of head rotation, the rate of change of angular velocity (degrees/s²), such that lower values indicate smoother movement. *Pause fraction* was defined as the time-weighted proportion of the phase during which head speed fell below a threshold. Separate thresholds were used for each phase since the distribution of head speed differs substantially between the two phases. Head speed during the follow phase was strongly bimodal, with the lower mode corresponding to physical head motion and the upper mode corresponding to motion of the virtual body. The two modes are separated by a near-empty interval, and the pause threshold was placed within it at 1.0 units/s. During the point phase, in which participants’ virtual body remained stationary and the head speed distribution was not bimodal, the pause threshold was set at the sample median of 0.02 units/s.

##### Path and duration ((2 x 2) + 2; 6 features)

Total path length traversed by the HMD (units) and total duration (s) were computed for both phases (4 features). The point phase was also subdivided into the interval from the endpoint cone to lighthouse identification and the interval between lighthouse and cabin identification (2 features), which together sum to the point-phase duration.

##### Controller movement (1 x 2; 2 features)

Mean movement speed of the right controller (units/s) was computed for both phases. Left controller data were excluded because participants used the left controller only to review task instructions.

##### Spatial error ((2 x 3 landmarks) + 2; 8 features)

Distance error (units) and angular pointing error (degrees) between the participant’s indicated position and the true position were recorded for the start location, lighthouse and cabin (6 features), together with the mean distance error and mean angular error across the three landmarks (2 features). These are properties of the pointing response and not computed per phase. These are similar to the error metrics (Sec 2.4.1), whereas the other 20 parameters were derived from HMD and controller data, and are termed *kinematic variables*.

All rate-based features were computed from true per-frame time intervals rather than assuming uniform frame spacing, as sampling was irregular. Duplicate frames, identified by identical timestamps and identical position and orientation values, were removed prior to any rate computation to avoid division-by-zero artifacts.

#### 2.6.2 SVM modeling

Trial-level features were aggregated at the participant level by taking the mean across each participant’s first 16 trials, giving one feature vector per participant. The primary model used the 20 kinematic features, that is, all groups in Sec. 2.6.1 except spatial error; a comparison model additionally included the 8 spatial-error features. A support vector classifier separates two classes by fitting the boundary that maximises the margin between them in a feature space induced by a kernel function. We used a radial basis function (RBF) kernel, 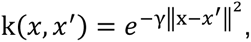 which measures similarity between pairs of feature vectors (*x*, *x*^′^) as a decaying function of distance. The kernel coefficient *γ* sets the width of the kernel: small *γ* gives each training point influence over a wide region of feature space. A regularisation parameter *C* sets the penalty for training points that fall inside the margin or on the wrong side of the boundary: small *C* tolerates more violations and yields a smoother boundary, large *C* fits the training participants more closely. Class weights were balanced by inverse group frequency, so that the 33 younger and 21 older participants contributed equally. Features were z-scored, scaled to 53 training participants and applied unchanged to the held-out participant. Predicted probabilities were obtained by Platt calibration, fitted within the training set of each fold.

Hyperparameters were selected by nested cross-validation. Within each outer fold, the model was tuned over a grid of five *C* values (1, 10, 50, 100, 200) crossed with six *γ* values (0.00001, 0.0001, 0.001, 0.01, 0.1, and scikit-learn “scale” default), giving 30 combinations. The “scale” default sets *γ* = 1/(*nVar*(X)), where *n* is the number of features and *Var*(X) the variance of the training matrix; for z-scored input (*Var*(X) = 1) this is 1/20 = 0.05 for the primary model and 1/28 = 0.036 for the comparison model. Each combination was scored by leave-one-participant-out cross-validation over the 53 training participants of that outer fold, so tuning never saw the held-out participant. Combinations whose inner AUC fell below 0.5 or could not be scored were discarded. The best-scoring combination was then refitted on all 53 training participants and used to classify the held-out participant. In total, this is 54 outer folds, each running 53 inner folds over 30 combinations.

BA is the mean of the two group recall rates, so moving one participant across the decision boundary changes it by 1/(2n), where n is the size of that participant’s own age group. In the inner folds this is ∼1.5% for a younger participant and ∼2.5% for an older one, so settings differing by less than that are not distinguishable. Combinations within 0.02 BA of the best, about one participant’s worth, were therefore treated as tied. Among tied combinations we chose the one with *C* closest to 1, breaking any further ties toward the smallest *γ*. Since *C* = 1 is the most strongly regularised setting in the grid, this prefers the simpler model over whichever setting happened to edge ahead on a coarse inner score.

Predicted probabilities of older-group membership were taken from the Platt-calibrated output, and class labels were assigned by the per-fold threshold that maximised the Youden index (Sec. 2.6). BA, sensitivity and specificity in Table 5 are computed from these labels, and AUC from the same calibrated probabilities.

#### 2.6.3 MiniRocket modeling

The raw time series was modeled with MiniRocket (Dempster et al., 2021), a deterministic variant of the ROCKET transform (Dempster et al., 2020), using the variable-length multivariate implementation (MiniRocketMultivariateVariable) in sktime (Löning et al., 2019). ROCKET-family methods convolve each input series with a large bank of fixed, untrained kernels and reduce every kernel response to a single pooled statistic. The classifier is then applied to the statistics across kernels.

MiniRocket uses a fixed bank of 84 distinct kernel patterns and 119 features per kernel. Each kernel is applied at up to 32 distinct dilations, spaced logarithmically with respect to series length, and several bias terms are drawn per dilation, from the quantiles of the convolution output computed on a randomly chosen training series. One feature, the proportion of positive values (PPV) of the convolution output after subtracting the bias, was computed per kernel, dilation and bias combination, giving 84 x 119 = 9,996 statistics per trial.

Input consisted of the 14 kinematic channels described in Sec. 2.4.2, taken from the follow phase only and from each participant’s first 16 trials. The transform accepts variable-length series, so trials were used at their native durations without resampling, truncation or padding, with dilations scaled relative to the longest series in each training set.

The transform was fitted on the training participants within each outer leave-one-participant-out fold and then applied to both the training trials and the held-out participant’s trials. Mean kernel features across trials were computed for each participant, z-scored across the training participants only, and classified with a radial basis function support vector classifier with class weights balanced by group frequency and predicted probabilities obtained by Platt calibration.

#### 2.6.4 Late fusion

The late fusion model combined the kinematic SVM with MiniRocket. For each participant, the two models’ predicted probabilities of belonging to the older group were averaged with equal weight, and the fused probability was labeled with the per-fold Youden threshold.

No model was refitted for fusion - each model had already produced a probability for every participant while that participant was held out, and fusion averaged these probabilities. Both inputs to each participant’s fused probability therefore came from models trained without that participant, and the threshold applied to it was chosen from the fused probabilities of the other 53 participants.

## 3 RESULTS

### 3.1 Sample Characteristics

The final sample comprised 54 adults aged 18 to 84 years (younger: < 55 years; n = 33, M = 27.21, SD = 11.59; older: ≥ 55 years; n = 21, M = 68.57, SD = 6.64). The two age groups differed significantly on several participant characteristics (Table 1). Older adults reported more years of education (19.19 vs. 16.27 years) and scored lower on the Corsi task (4.90 vs. 6.09), and PMAT (15.90 vs. 18.91), and had lower self-reported gaming (0.71 vs. 1.79) and VR experience (0.38 vs. 0.97). Older adults also reported lower state anxiety (5.43 vs. 6.33), trait anxiety (7.14 vs. 9.45), and depressive symptoms (3.38 vs. 8.82). Age was correlated with every continuous covariate (all Spearman |ρ| ≥ .35): education (ρ = .66), depressive symptoms (ρ = −.53), gaming experience (ρ = −.50), VR experience (ρ = −.48), Corsi span (ρ = −.45), trait anxiety (ρ = −.43), state anxiety (ρ = −.38), and PMAT (ρ = −.35) (Table S12). All older adults were cognitively screened using the MoCA (M = 28.76, SD = 1.34, range 26-30).

### 3.2 Error Metrics Results

#### 3.2.1 Attrition

49/54 participants completed all 30 trials (32/33 younger, 17/21 older). Of the five non-completers, all withdrawals occurred for reasons unrelated to task performance: four for cybersickness or dizziness and one due to time constraints. Older non-completers had lower Start Error in blocks 1-3 than older completers (29.1° vs. 59.5°) but higher error on the four remaining metrics (Table S1). Removing these participants should therefore inflate the Start Error age effect and attenuate the others, which is confirmed by the completers-only sensitivity analysis (Table S6).

#### 3.2.2 Error Magnitudes by Age Group

Participant-level mean absolute errors are shown in Table 2. Older adults showed larger errors than younger adults on 4/5 metrics, with the largest difference on Start Error (52.2° vs. 35.6°, Hedges’ g = 0.64, 95% CI [0.08, 1.19]). Differences were smaller for Lighthouse Error (g = 0.45), Start-LH Error (g = 0.43), and Cabin Error (g = 0.41). LH-Cabin Error ran in the opposite direction, with younger adults showing larger errors than older adults (48.3° vs. 42.1°, g = −0.20). These descriptive comparisons are uncorrected for multiplicity and are not statistically independent of the continuous-age models reported below.

**Table 2.** Observed and Model-Estimated Error Magnitudes by Age Group.

| Metric | Younger M (SD) | Older M (SD) | Diff | Hedges g | 95% CI | Fitted at 27 | Fitted at 69 |
|---|---|---|---|---|---|---|---|
| Start Error | 35.6 (24.4) | 52.2 (26.9) | +16.6 | 0.64 | [0.08, 1.19] | 30.6 [25.8, 36.3] | 47.6 [38.7, 58.5] |
| LH Error | 55.9 (20.0) | 65.3 (22.0) | +9.4 | 0.45 | [-0.10, 0.99] | 52.7 [46.2, 60.1] | 61.3 [52.2, 71.9] |
| Cabin Error | 58.7 (21.4) | 67.7 (21.9) | +9.0 | 0.41 | [-0.14, 0.95] | 55.3 [47.7, 64.0] | 62.0 [51.9, 74.1] |
| Start-LH Error | 44.3 (14.6) | 50.9 (16.7) | +6.6 | 0.43 | [-0.12, 0.97] | 42.6 [37.9, 47.8] | 48.3 [41.9, 55.7] |
| LH-Cabin Error | 48.3 (30.8) | 42.1 (29.8) | -6.2 | -0.20 | [-0.74, 0.34] | 40.9 [33.6, 49.7] | 33.8 [26.5, 43.1] |

Note. Observed values are participant-level means across blocks 1-6, in degrees. Positive Hedges' g indicates larger error in older adults. Fitted values are estimated marginal means from the continuous-age GLMM (Table 3), evaluated at ages 27 and 69 (the observed group mean ages) at the session midpoint (block\_c = 0), with 95% confidence intervals.

The five error magnitudes were not statistically independent of one another (Table S11a). Start-LH Error and LH Error magnitudes were near-redundant (Spearman ρ = .97), as expected given their shared construction, and LH and Cabin Error magnitudes were also strongly correlated (ρ = .93). Start Error was comparatively independent of the landmark metrics (ρ = .14 to .22) and negatively related to LH-Cabin Error (ρ = −.28). The five metrics therefore reflect approximately three independent sources of variance.

#### 3.2.3 GLMM Model Specification

A Gamma distribution fitted all five error metrics better than a lognormal distribution (ΔAIC = 261 to 415 in favour of Gamma; Table S2a). Intraclass correlations from intercept-only models ranged from .13 to .36 (Table S2b), confirming that a by-participant random intercept was warranted while indicating that most variance in trial-level error was within rather than between participants. Likelihood ratio tests (Table S2c) supported age-dependent dispersion for Start Error (*χ*²(1) = 25.52, *p* < .001), Cabin Error (*χ*²(1) = 17.70, *p* < .001), and Lighthouse Error (χ²(1) = 8.24, p = .004) magnitudes, but not for Start-LH Error (*χ*²(1) = 2.41, *p* = .120) or LH-Cabin Error (*χ*²(1) = 1.00, *p* = .318). All models converged with finite standard errors.

#### 3.2.4 GLMM analysis

##### Age

The age effects for each metric are outlined in Table 3. Start Error magnitude increased by 1.06% per year, corresponding to a 54% larger error across the 41.36-year gap between the two groups’ mean ages (ratio = 1.54, p < .001, Holm p = .004). No other error magnitudes showed a reliable age effect: LH Error (ratio = 1.16, Holm p = .544), Cabin Error (ratio = 1.12, Holm p = .544), Start-LH Error (ratio = 1.13, Holm p = .544), and LH-Cabin Error (ratio = 0.83, Holm p = .544).

**Table 3.** Error Metrics Analysis by Continuous-Age GLMM.

| **Metric** | **% per year** | **Ratio** | **95% CI** | **p** | **p Holm** |
|---|---|---|---|---|---|
| Start Error | +1.06 | 1.54 | [1.20, 1.99] | < .001 | .004 |
| Lighthouse (LH) Error | +0.36 | 1.16 | [0.95, 1.41] | .136 | .544 |
| Cabin Error | +0.27 | 1.12 | [0.90, 1.39] | .307 | .544 |
| Start-LH Error | +0.30 | 1.13 | [0.95, 1.35] | .157 | .544 |
| LH-Cabin Error | -0.45 | 0.83 | [0.62, 1.11] | .212 | .544 |

Note. N = 54 participants, 1,535 trials across blocks 1-6. Gamma GLMM with log link, by-participant random intercept, and age-dependent dispersion. Ratios and confidence intervals rescale the per-year coefficient across the observed 41.36-year gap between group mean ages. Age ratio of 1.00 indicates no difference. p Holm reflects Holm correction within this five-outcome family. Block and interaction terms are reported in Table S3.

##### Block

Error magnitudes decreased across blocks for LH-Cabin Error (−7.0% per block, Holm p < .001) and Cabin Error (−4.1% per block, Holm p = .010). LH Error showed a trend that did not survive correction (−3.1% per block, Holm p = .081). Start Error and Start-LH Error magnitudes did not change significantly across blocks (both Holm p > .99) (Table S3).

##### Age x Block

No interaction survived correction. The largest effect was for LH-Cabin Error, where the error magnitude reduction across the session was attenuated with increasing age (ratio across the group gap = 1.06, p = .020, Holm p = .102) (Table S3).

#### 3.2.5 Trial-to-Trial Variability in Error Magnitude

Trial-to-trial variability was greater in older than younger adults for Start Error (mean participant-level std 35.5° vs. 28.2°, Hedges’ g = 0.59, *p = .*036). However, this difference did not survive Holm correction across the five metrics (Holm *p* = .178). No other metric reached significance. Differences were 2.47° for the Lighthouse Error (g = 0.25, Holm *p* =.795), 0.75° for the Cabin Error (g = 0.07, Holm *p* =.809), and 3.01° for the Start-LH Error (g = 0.31, Holm *p* =.795). LH-Cabin Error differed in the opposite direction, with 4.73° (14.1%) lower variability in older adults (g = -0.50; Holm *p* = .301) (Table 4).

**Table 4.** Age-Related Differences in Trial-to-Trial Variability.

| **Metric** | **Younger SD** | **Older SD** | **Difference [95% CI]** | **Hedges g** | ***p*** | ***p* Holm** |
|---|---|---|---|---|---|---|
| Start Error | 28.2 | 35.5 | 7.23 [0.64, 13.68] | 0.59 | .036 | .178 |
| Lighthouse (LH) Error | 45.4 | 47.9 | 2.47 [-2.40, 7.51] | 0.25 | .376 | .795 |
| Cabin Error | 46.4 | 47.2 | 0.75 [-5.24, 6.18] | 0.07 | .809 | .809 |
| Start-LH Error | 37.9 | 40.9 | 3.01 [-1.92, 7.98] | 0.31 | .265 | .795 |
| LH-Cabin Error | 33.4 | 28.7 | -4.73 [-9.80, 0.53] | -0.50 | .075 | .301 |

Note. Trial-to-trial variability is defined as the standard deviation of absolute error across each participant's trials in blocks 1-6, in degrees; the presented values are group means of per-participant standard deviations. A positive difference and a positive Hedges' g indicate greater variability in older adults. P-values are from two-sided Monte Carlo permutation tests (100,000 resamples) randomly re-assigning participants to age groups; confidence intervals are percentile bootstrap (10,000 resamples). p Holm reflects Holm correction across the five metrics. Mean error magnitudes for the same participants are reported in Table 2. Because the standard deviation is computed across all six blocks, it incorporates within-session change as well as trial-to-trial noise. The completers-only sensitivity analysis is reported in Table S5.

This pattern was unchanged in the balanced completers-only panel. The Start Error difference was larger (9.34°, 32.7% greater variability; g = 0.77; *p* = .013; Holm *p* = .064), and the remaining four metrics again showed no significant difference (all *p ≥ .160 and* Holm *p* ≥ .639; Table S5). The correlation between the number of trials a participant contributed and their standard deviation was negligible for every metric (Spearman ρ = −.09 to .06 across the five metrics; Start Error ρ = -.01).

### 3.3 Machine learning results

The error metrics analyses characterise how accurately participants pointed. The ML analyses ask whether the movement produced during the task carries information about age group. Data was restricted to each participant’s first 16 trials and all models were evaluated by leave-one-participant-out cross-validation.

#### 3.3.1 Model comparison

Four models were compared (Table 5). The SVM trained on all 28 features, with hyperparameters selected by nested cross-validation, performed at 73.4% BA (AUC 0.772). Removing the 8 task-performance features derived from the pointing responses and retaining only the 20 aggregated kinematic features increased accuracy to 80.7% BA (AUC 0.854), correctly classifying 25/33 younger and 18/21 older adults. MiniRocket, applied directly to the 14 raw trajectory channels with no feature engineering, reached 65.2% BA (AUC 0.827). Late fusion of the kinematic SVM and MiniRocket, combining the feature-based and time-series decision values within the cross-validation loop, gave the highest balanced accuracy (82.0%, AUC 0.860), correctly classifying 29/33 younger and 16/21 older adults.

**Table 5.**
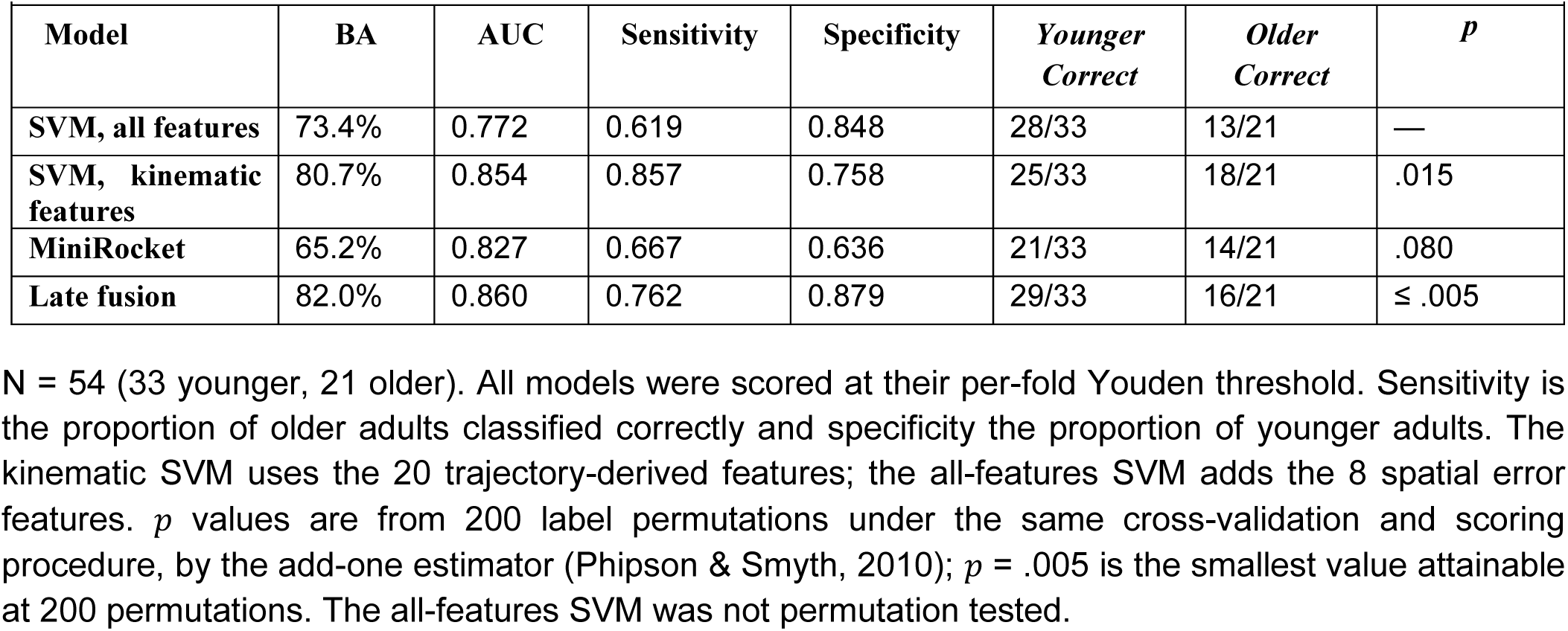
Classification performance by model.

The fusion model’s advantage over the kinematic SVM alone is small: 1.3% BA and 0.006 AUC. At n = 54 a single participant is worth 1.5% BA in the younger group and 2.4% in the older, so fusion gains less than one participant’s worth. The two models also differ in where they are accurate rather than in how accurate they are. Fusion classifies four more younger adults correctly but two fewer older adults, so the kinematic SVM alone is the more sensitive of the two. MiniRocket ranked participants well (AUC 0.827) but classified comparatively few correctly at its operating point (BA 65.2%), and its thresholds varied widely across folds (0.28 to 0.49), so the information in its ranking did not carry through to the thresholded labels.

The ROC curves (Fig. 4) show the same, with the fusion model and the kinematic SVM close together across most of the range and MiniRocket lying below them at low false positive rates. The confusion matrices (Fig. 5) show the per-model accuracy: the kinematic SVM classifies 25/33 younger and 18/21 older adults correctly, MiniRocket 21 and 14, and fusion 29 and 16.

**Figure 4.**
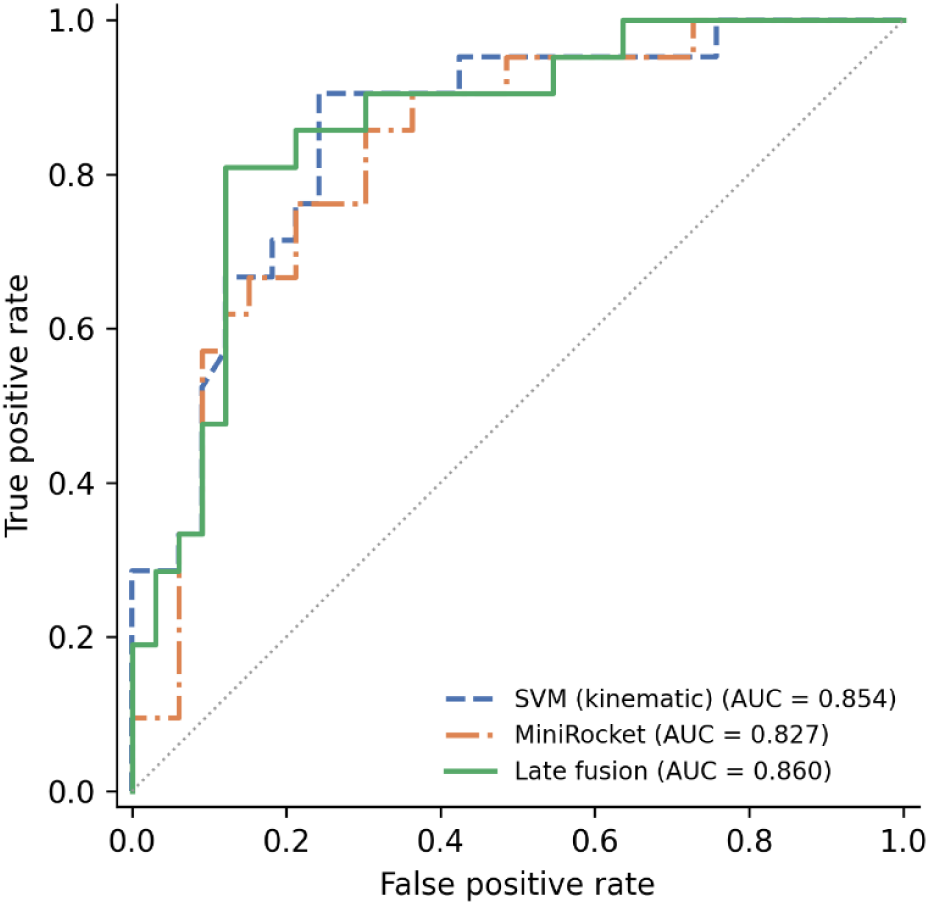
Receiver operating characteristic (ROC) curves for age group classification using three ML models. Curves are derived from held-out predicted probabilities pooled across all 54 leave-one-participant-out folds. The diagonal indicates chance performance. AUC valued given in the legend correspond to those reported in Table 5.

**Figure 5.**
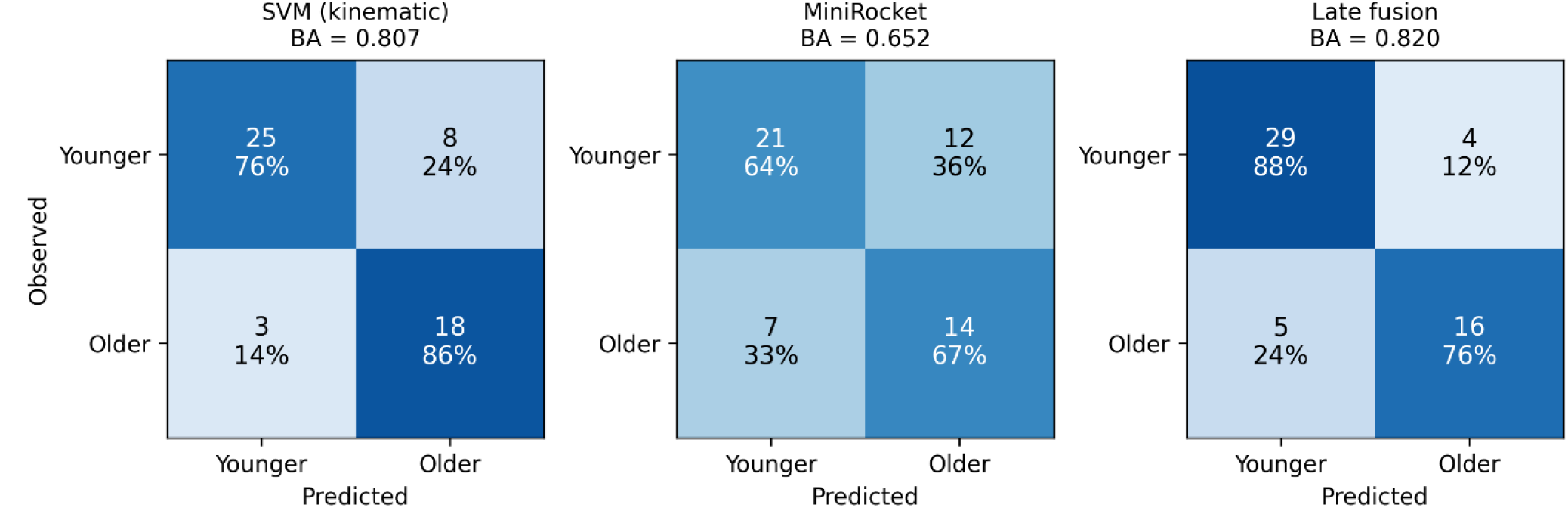
Confusion matrices for the three classification models. Rows are true age group and columns are predicted age group; cells are shaded by row proportion and annotated with the number of participants. N=54 (33 younger, 21 older). Balanced accuracy (BA) is given above each panel.

#### 3.3.2 Permutation test

To test whether the observed BAs could arise by chance under the leave-one-participant-out scheme, age-group labels were randomly permuted across participants and the complete pipeline was rerun under each permutation. This included both component models, with nested hyperparameter selection for the SVM repeated inside every outer fold, and the per-fold Youden threshold. The MiniRocket transform and the z-scoring of its output features do not depend on labels, so those per-fold matrices were computed once and reused across permutations, while the classifier fitted to them was refitted under each permuted label vector. Since both component models were recomputed under each permutation, the kinematic SVM, MiniRocket and their fusion were all scored against the same set of permuted labels, giving a null distribution for each at no additional cost.

Two hundred permutations were run, each with a fixed and recorded seed, so the null distribution can be regenerated exactly. The p-value was computed as (*E* + 1)/(*B* + 1), where *E* is the number of permutations reaching the observed BA and *B* the number of permutations, which gives a smallest attainable p-value of 1/201 = .005 (Phipson & Smyth, 2010).

The BA of the kinematic SVM, 80.7%, was exceeded by 2/200 permuted runs (p = .015), against a null centred on chance (mean 52.2%, SD 8.7%, maximum 84.4%; Supp. Fig. 2). The fusion model BA of 82.0% was exceeded by none (p ≤ .005; null mean 49.9%, SD 8.7%, maximum 81.4%). MiniRocket BA of 65.2% was exceeded by 15/200 (p = .080; null mean 51.1%, SD 9.7%, maximum 84.4%), so its BA is not distinguishable from chance even though its ranking of participants is informative (AUC 0.827). This is the same threshold instability described in Section 3.3.1.

The feature subset was held fixed across permutations while hyperparameters were re-selected. These tests therefore address whether each model exceeds chance, and not whether the kinematic subset outperforms the full feature set, a comparison that was made on held-out folds and is treated as exploratory (Section 4.8).

## 4.0 DISCUSSION

We ran a navigation study where participants across the adult age spectrum traversed a virtual maze, and in each trial, pointed back to their start location and the location of two landmarks that were hidden from them during the traversal. Navigation involves several interconnected neural computations, and we designed this task so that participants could engage these processes at various phases of the trial. Keeping track of one’s own position in the absence of landmarks and then reporting the start position involves path integration, locating the landmarks after revealing the true start location involves reorientation, and locating one landmark based on the other involves allocentric memory. We found that (1) age effects are expressed in path-integration-based metrics and not so much after reorientation, and (2) the movement during the traversal is more informative of age than accuracy measures. In summary, age effects are prominent in how you move during episodes of path integration, more so than how well you performed.

### 4.1 Age effects are reliably expressed in path integration

Age was reliably associated with Start Error, which involved pointing back to the start location after traversing with no visible external cues, a measure of path integration. Start Error increased by 1.06% per year. Four other metrics (Lighthouse, Cabin, Start-LH and LH-Cabin Error), all of which were based on responses after external cues had been restored (hereafter referred to as post-reorientation metrics), showed no reliable age effect after corrections. This finding is consistent with literature suggesting that path integration is an age-sensitive component of human navigation (Lester et al., 2017; Segen et al., 2022; Stangl et al., 2020). A possible mechanistic explanation is that ageing makes the movement information used for path integration noisier, rather than causing failure of path-integration itself (Stangl et al., 2020). Individual differences in entorhinal grid-cell-like representations have been shown to predict path integration error in older adults (Stangl et al., 2018), providing a candidate neural substrate for the observed behavioural gradient.

The Start Error effect did not depend on how age was specified or on attrition. Replacing continuous age with the binary younger/older classification produced the same pattern. Restricting the sample to the 49 participants who completed all 30 trials strengthened the estimate. The four older adults who withdrew had lower Start Error before withdrawal than those who continued to the end (Table S1). Adding sex, education and Corsi span (a measure of visuospatial working memory) attenuated the error increase from 54% to 47%, albeit with a confidence interval range statistically distinguishable from no age difference. Education and Corsi span were themselves strongly correlated with age in this sample (ρ = .66 and -.45). Adjusting for them therefore estimates the age difference between people of equal education and working-memory span, which removes any part of the age effect that may operate through these characteristics. We therefore treat the adjusted estimates as sensitivity analyses rather than as corrected estimates of the total age effect.

Older age was associated with 26-64% larger Start Error, and the effect never reversed in direction (Tables S6-S8, S18). However, the certainty of the estimate varied: statistical support weakened as age-correlated characteristics were added, and after adjustment for gaming experience the confidence interval included 1.00, indicating that the age effect was no longer statistically distinguishable from no association. Gaming experience, education and Corsi span all differ with age in this sample, and therefore these data establish a consistent age-associated difference in path integration error but cannot reliably determine whether it is independent of these characteristics.

If older adults’ larger Start Error reflected difficulty with the apparatus or interface, the difference between groups should decrease as participants gain experience with the task. However, it does not: Start Error showed no change across the six blocks and no age x block interaction. Meanwhile the metrics that did improve across the session – Cabin Error and LH-Cabin Error – are the ones that showed no age effect. Adjustment for VR experience specifically, which is the more direct index of interface familiarity in the present task, produced only a modest attenuation of the age estimate (1.47 to 1.40), whereas adjustment for gaming experience produced a larger one (1.47 to 1.26). Because path complexity also increased across blocks, stable Start Error could reflect improvement that is offset by harder paths. The data therefore argue against simple learning effects, but do not rule out a contribution of prior technology experience.

### 4.2 Null age effects on post-reorientation metrics

The absence of a reliable age effect on Lighthouse, Cabin, Start-LH and LH-Cabin Error should not be read as evidence that the corresponding processes are preserved with age. The confidence intervals remain wide – Cabin Error, for example, is estimated at ratio = 1.12, 95% CI [0.90, 1.39] – and is compatible with as much as a 39% increase in older adults. We instead suggest that the age signal in this paradigm is concentrated in the phase where participants must rely on self-motion alone and is not detectable in the phases where external cues have been restored.

All four metrics with null effects were calculated from responses made with the true start position and the sun visible, and these cues may provide an opportunity to recalibrate the accumulated path-integration estimate (Epstein et al., 2017). Restoring a reliable global anchor before the landmark responses may reduce group differences on those responses. This is consistent with evidence that older adults retain flexible use of spatial reference frames and effective strategy selection even where representational precision is reduced (Ladyka-Wojcik & Barense, 2021; McAvan et al., 2021), and that age differences in spatial memory are attenuated under more naturalistic, cue-supported navigation (Hill et al., 2024).

The landmark metrics also differ in how much stable between-participant variance they contain. Intraclass correlations from the intercept-only models were highest for Start Error (.312) compared to the post-reorientation metrics (Lighthouse .180; Cabin .209; Table S2b). These low values indicate that the post-reorientation measures contained proportionally more within-individual variation and less stable between-individual variation, which may reduce the precision with which participant-level predictors such as age can be estimated.

### 4.3 Trial-to-trial variability and the noise-accumulation account

Start Error magnitude was the only metric in which older adults were more variable trial-to-trial. Their standard deviations were 25.6% larger than those of younger adults (Table 4). Lighthouse, Cabin and Start-LH Error variability showed differences in the same direction but of negligible size (1.6% to 7.9%), while LH-Cabin Error differed in the opposite direction, with 14.1% lower variability in older adults. Neither the Start Error difference nor any other metrics reached significance after Holm correction in either the full sample or the completers-only panel; we therefore treat these results as suggestive.

The direction and selectivity of the variability differences are nonetheless consistent with the noise-accumulation account proposed by (Stangl et al., 2020). Given a noisier velocity signal, the responses that depend on integrating self-motion should become less consistent and less accurate from trial to trial. Responses made with external cues available can be anchored to environmental features and should therefore be less affected by noise in that estimate. This is the pattern that was observed: Start Error is the only response that is produced without visible external cues, and it was the only response where variability differed appreciably with age.

The LH-Cabin variability was the only one that showed a difference in the opposite direction. As a relational metric, LH-Cabin Error captures the angular separation between the two landmark responses rather than the accuracy of either relative to its target. Its trial-to-trial variability could therefore arise from either response varying, or both, in ways the metric cannot distinguish.

A number of factors might affect the variability analysis. Since the standard deviation was computed across all six blocks, it incorporates systematic within-session change and could therefore be inflated. However, this trend confound does not affect the Start Error comparison, which showed no significant change across blocks in either group (Sec. 4.5), suggesting that the standard deviations reflect trial-to-trial variability rather than session-long drift. Absolute angular error is bounded at 0° and 180°, so spread is necessarily compressed as the mean approaches the middle of this range. Since older adults had substantially higher mean error, some compression of their variability estimates is expected. Variability was also computed on absolute rather than the signed errors, which discards the direction of each deviation. The resulting measure therefore captures only how much the size of the deviation varied, and not how widely responses scattered around the true bearing. A circular measure of dispersion computed on signed responses would be an appropriate approach in a sample large enough to support participant-level variance comparisons with adequate power.

### 4.4 Distinct computational processes during navigational phases

Does our task distinguish distinct computational processes, as intended? Between participants, the intercorrelations among metric magnitudes (Table S11a) show one dimension that is essentially independent of all others (Start Error), one post-reorientation landmark dimension (Lighthouse, Start-LH and Cabin), and a relational dimension that is negatively related to Start Error (LH-Cabin, ρ = −.28).

Within trials, it is expected that the lighthouse and cabin responses are coupled, as the placed lighthouse marker remained visible while pointing to the cabin, which is exactly what was observed (directional r = 0.26, Holm p < .001; magnitude r = 0.21, Holm p = .002). Conversely, if the reorientation stage successfully re-anchors the participant to an external frame, the start and landmark responses should be decoupled, which is also what was observed (Start-Lighthouse directional r = −0.006; Start-Cabin r = −0.022). The absence of detectable Start–Lighthouse and Start–Cabin coupling suggests that individual differences in the initial path-integration response were not carried forward into subsequent landmark pointing.

Consistent with this logic, participants who spontaneously described using the sun as a directional cue showed weaker Start-Lighthouse directional coupling than those who did not (difference = −0.146, p = .031). The sun was visible as a global directional cue only after the start response and before the landmark responses, so weaker coupling is expected in participants who used it. However, the effect did not survive correction (Holm p = .093), rests on 15 participants, and depends on a classification drawn from unprompted responses; it is reported anecdotally. Future designs should consider randomizing sun availability and using structured questions on navigational strategy rather than an open-ended description.

### 4.5 Session position, learning, and the age x block interaction

The experiment was divided into 6 increasing ‘complexity’ blocks of 5 trials each. Error magnitudes decreased across blocks for LH-Cabin Error (−7.0% per block, Holm p < .001) and Cabin Error (−4.1% per block, Holm p = .010), with a comparable trend for LH Error that did not survive correction (−3.1% per block, Holm p = .081). Start Error and Start-LH Error did not change (both Holm p > .99). Since path complexity increased with every block, a change across blocks could reflect practice, the specific path configurations of later blocks, or both. The pattern of block effects is informative, nonetheless. Error magnitudes only decreased reliably for Cabin and LH-Cabin Error, with a weaker trend for Lighthouse Error. These responses depend on the lighthouse and cabin, whose positions were fixed throughout the task and could therefore be learned across trials. In contrast, Start Error and Start-LH Error – which are both referenced to a start location that changed on every trial, requiring novel path integration computations – showed no detectable change. This dissociation suggests participants learned the fixed allocentric landmark layout over the session.

Only one age × block interaction approached significance, on LH-Cabin Error, and was not stable across age specifications (continuous: ratio = 1.06, Holm p = .102; binary: ratio = 1.11, Holm p = .001). The descriptive pattern (Table S14a) shows that age-based deterioration is not what drives this effect: younger adults began the session with substantially larger LH-Cabin Error than older adults (60.7° versus 38.6° in block 1) and converged by the end (41.1° versus 41.4° in block 6). The interaction therefore reflects greater change in the younger group from a worse starting point at the beginning of the task, rather than a widening age gap. Both groups also showed a large and highly consistent negative directional bias on this metric, underestimating the true angular separation between the two landmarks by roughly 32° to 38° (mean resultant lengths .82 to .86; Table S13), which suggests that LH-Cabin Error is consistent with a systematic compression of perceived angular separation shared across age groups.

To isolate practice effects, we introduced an additional block 7 as a repetition of block 1, later during data collection. Although the results point numerically towards learning on all five metrics (changes of −9.8° to −23.0°, Table S15b), this data is only available from 13 participants, and no change survived correction (smallest Holm p = .109). Counterbalanced or interleaved block order may resolve this in a future iteration of the paradigm.

### 4.6 Kinematics outperform error metrics

Movement kinematics discriminated age group across the sample without depending on any single pointing response. The SVM trained on the 20 kinematic features reached 80.7% BA and the late fusion of that model with MiniRocket reached 82.0%, both exceeding chance under label permutation (p = .015 and p ≤ .005). MiniRocket alone (65.2% BA) was not distinguishable from chance (p = .080), although its ranking of participants was informative (AUC 0.827). The kinematic SVM is more sensitive than specific (0.857 against 0.758), so older adults are classified more reliably than younger adults; the fusion model reverses this (0.762 against 0.879), gaining younger adults and losing older ones. With 54 participants, differences of the size separating adjacent rows in Table 5 correspond to a small number of individuals, and the ordering of models should be read as indicative rather than as a precise ranking.

These classification results complement rather than replace the error-metric analyses in Section 3.2. The classification analyses ask how much age-related information is recoverable from movement without specifying in advance which aspect of movement carries it. The two findings are consistent: the movement produced during navigation and recall appears to contain age-related structure that per-trial summary statistics do not fully capture, including the accuracy-based measures that form the basis of the error-metric models.

The two sources of information were compared directly by splitting the 28 features into the 20 derived from movement and the 8 derived from the pointing responses, and training a classifier on each under the same leave-one-participant-out structure. The movement features gave 80.7% BA and 0.854 AUC. The task-performance features alone gave 0.483 AUC, which is at chance, so no age-group information was recoverable from them in this form. This supports the claim that continuous movement carries age-related information that these discrete error metrics do not. A movement-based classifier can be more naturalistic, and requires no response the participant knows is being evaluated.

### 4.7 Complementarity between feature-based and time-series models

The MiniRocket model is trained on continuous time series of movement, while the SVM is trained on aggregated hand-crafted features of movement. The two models differ sharply in how stable their operating points are. The thresholds MiniRocket selects vary widely across folds (0.28 to 0.49), while the kinematic SVM’s vary hardly at all (0.379 to 0.384). An unstable threshold means the operating point chosen on one set of participants transfers poorly to the next, which is why MiniRocket’s BA was not distinguishable from chance under permutation (p = .080) even though its ranking of participants carries real age-related information. The two models also make different mistakes. Against the 33 younger participants, the kinematic SVM produced 8 false positives and MiniRocket 12, overlapping on 5. Three of the kinematic SVM errors and seven of the MiniRocket errors are unique, so most of MiniRocket’s younger-group error is unique to it. The fusion model that combines them exceeds both in BA and AUC, but not in sensitivity: the kinematic SVM alone identifies 18/21 older adults against the fusion model’s 16. Fusion’s advantage is a net reduction in errors across the sample rather than the rescue of the cases that defeated both models. Probability averaging under a threshold selected inside the fold moves participants across the decision boundary according to how confident each model was, not according to whether the models agreed. SVM features are computed separately for the follow and point phases, while MiniRocket’s convolutional kernels operate over the raw sequence without phase labelling. Whether this difference explains why the models make different mistakes was not tested.

### 4.8 Limitations

#### Sample size and composition

We analyzed 54 participants in a 33/21 group split. BA and AUC estimated by leave-one-participant-out cross-validation at this size are imprecise, and differences of a few points between models correspond to a small number of participants changing classification. The younger group is drawn substantially from the university subject pool, so age is confounded with student status. The older group was screened for normal cognition (MoCA ≥ 26) while the younger group was not.

#### Age specification

The binary split at 55 years reflects the observed bimodal age distribution rather than a theoretical boundary, and was chosen after inspecting the data. Middle age is sparsely represented, so the analysis cannot speak to when or how gradually the differences it detects emerge. The continuous-age GLMMs address this limitation somewhat for the error metrics.

#### Attrition is age-asymmetric

Session length was shorter on average among older participants, which is why the classification analysis was capped at 16 trials. However, this means the models were not trained on the later, longer trials where group differences might be larger or smaller.

#### Interface familiarity

Older adults reported less gaming and VR experience than younger adults. The error-metric analysis found that adjusting for gaming experience attenuated the Start Error age effect below the conventional threshold, while adjusting for VR experience did not. The classification models were not adjusted for either. Whether performance is partly recoverable from interface familiarity rather than navigation ability is therefore untested and should be addressed directly, ideally by comparing experienced and inexperienced participants within the younger group.

### 4.9 Conclusion

Our study underscores the importance of a particular feature in the analysis of navigational processes relevant to aging – movement while path integrating. If you are keeping track of your own movement with respect to a start point, or tracking the position of hidden external landmarks, you are path integrating, i.e. adding up your noisy, multisensory movement inputs to build an imperfect estimate of where you might be. Your movement during this challenging cognitive process seems to change as you age. These movement signals are a potential behavioural correlate of the degradation of the update of egocentric and allocentric representations of self and environment in the brain’s hippocampal formation. These are the brain regions first affected in the canonical spread of Alzheimer’s disease (AD), and thus it stands to reason that movement patterns while path integrating would be different between individuals with AD and age-matched controls. Further work using machine learning analysis of continuous movement data needs to be done on these clinical populations, and on animal models of AD to establish the mechanistic link. However, we must be careful to distinguish motor correlates of degraded path integration from AD-induced motor deficits. A significant effect of early AD or other dementia subtypes on movement patterns during path integration, will increase the efficacy of longitudinal, multimodal behavioural testing of dementia-related cognitive decline in older individuals.

## DATA AND CODE AVAILABILITY

Anonymized data can be made available upon reasonable request, in consultation with the UBC Office of Research Ethics.

Code for discrete event data analysis is available at: https://github.com/NC4Lab/kim2026movement_analysis_eventdata

## ACKNOWLEDGEMENTS

We thank all participants for their time and contribution to this study. We are grateful to former lab manager Isaac Morgan for his feedback on task design and support with obtaining behavioural research ethics approval and project administration, and to research assistants Talia Apel and Leticia Cid for data collection. We thank Han Cat Nguyen for helping to develop the initial version of the VR task software, the UBC Emerging Media Lab staff (Frederik Kjaer Svendsen, Mohsen Movahedi), student team (Mariane Olivan, Eric Leung) and leadership (Saeed Dyanatkar, Maryann Kempthorne) for developing it into a functional application. We thank Inzaghi Moniaga for implementing major design changes and fixing software bugs, and Amr Sadek and Matin Narimani for final refinements of the software. We are further grateful to Dr. Daniela Palombo who advised on the neuropsychological assessment battery and to Dr. Thalia Field for her feedback as a clinical co-investigator. Amy Wong designed Fig. 1(a) from an image generated by Claude AI.

## AUTHOR CONTRIBUTIONS

A.K. performed and supervised design and development of the task, data collection, and error metrics analysis. G.K. developed the ML models and analyzed data. A.W.L. and A.G.M. were instrumental in conceptualizing and designing the task and error metrics pipeline. A.G.M. co-developed and piloted the initial version of the task, and presented the methodology. M.S.M. supervised all trainees and all aspects of design, data collection and analysis. A.K., G.K. and M.S.M. drafted the manuscript and A.G.M. and A.W.L. edited and offered feedback.

## 1. SUPPLEMENTARY MATERIAL

## Model Diagnostics

Simulation-based residual diagnostics indicated departures from the Gamma model for all five error metric magnitudes under both age specifications (Table S9). Kolmogorov-Smirnov tests of uniformity were significant for every outcome (p < 0.001 to .030; Table S9a), as well as the dispersion tests (p ≤ .024; Table S9a). Dispersion statistics ranged from 0.64 to 0.84, indicating lower residual variability than expected under the fitted Gamma distributions. Participant-aggregated uniformity tests showed the same pattern (p < 0.001 to .038; Table S9b). Quantile deviation tests were significant for the Lighthouse and Cabin Error magnitudes, and for Start-LH Error under the binary specification only (Table S9a).

One possible explanation is that absolute angular error cannot exceed 180°, whereas the Gamma distribution has unbounded support. Thus, the fitted Gamma distributions may assign greater probability to extreme upper-tail values than is possible in the dataset. Residuals showed no systematic monotonic drift across blocks and comparable spread in the two age groups (median scaled residuals .49 to .57; interquartile ranges .46 to .56 against a target of .50; Table S9c), indicating that the misfit is likely distributional.

## Robustness to Age Specification (Age Group)

Replacing continuous age for the binary younger/older classification produced the same pattern of results (Table S4). Older adults showed 52% larger Start Error magnitude than younger adults at the middle of the session (ratio = 1.52, 95% CI [1.14, 2.02], p = .004, Holm p = .021), corresponding to estimated marginal means of 30.9° (95% CI [25.8, 37.0]) in the younger group and 46.9° (95% CI [37.5, 58.6]) in the older. No other age effect approached significance (all Holm ps = .661). The block effects were likewise unchanged (Cabin Error −4.9% per block, Holm p = .020; LH-Cabin Error −10.7% per block, Holm p < .001). Under this specification, the Age Group × Block interaction for LH-Cabin Error reached significance (ratio = 1.11, p < .001, Holm p = .001), indicating that younger adults improved across the session on this metric while older adults did not. Because this interaction did not survive correction under the primary continuous-age specification (Holm p = .102), it is not treated as a reliable effect and is noted only for completeness.

## Sensitivity Analyses

### Completers Only

Restricting the analysis to the 49 participants who completed all 30 trials strengthened the Start Error magnitude age effect in both age specifications (continuous: 1.20% per year; ratio across the group gap = 1.64, 95% CI [1.25, 2.15], p < .001; binary: ratio = 1.63, 95% CI [1.19, 2.23], p = .002). The four remaining effects were attenuated and the direction of all effects was preserved (Table S6).

### Covariate Adjustment

Adding sex, education, and Corsi span attenuated the Start Error magnitude age effect in both specifications, though the confidence intervals continued to exclude 1.00. (continuous: 0.93% per year; ratio across the group gap = 1.47, 95% CI [1.09, 1.97], p = .011; binary: ratio = 1.38, 95% CI [1.01, 1.89], p = .044). Holm correction across the five metrics within the adjusted models raises these values to p = .055 and p = .221, respectively (Table S7a). No individual covariate was reliably associated with any error metric after false discovery rate correction (all *q* ≥ .65; Table S7b). The variance inflation factors (VIF) were at or below 1.49 for all covariates in both adjusted specifications, indicating no collinearity problem (Table S10). Both PMAT and Corsi span are negatively correlated with age in this sample (ρ = -.35 and -.45, respectively) and are plausibly mediators of the relationship between age and navigation performance than confounds of it. Corsi span was selected as the single cognitive covariate in the adjusted models as it indexes the specific visuospatial working memory demands of the present task, whereas PMAT indexes general fluid reasoning.

### Interface Experience

Adding gaming experience to the adjusted model further attenuated the Start Error magnitude age effect (continuous: ratio = 1.47 to 1.26, p = .157; binary: ratio = 1.38 to 1.21, p = .234). Greater gaming experience was itself associated with smaller Start Error magnitudes (ratio = 0.82 per SD, p = .029), larger Cabin Error magnitudes (ratio = 1.17, p = .039), and larger LH-Cabin Error magnitudes (ratio = 1.28, p = .016), however neither survived correction across the pooled 10-test family (both q = .130). Adding VR experience attenuated the age effect less (continuous: ratio = 1.47 to 1.40, p = .032; binary: ratio 1.38 to 1.30, p = .123), and VR experience itself was not associated with any metric (all q ≥ .16). The age effect therefore remained nominally significant after adjustment for VR experience but not after adjustment for gaming experience. Adjusting for gaming experience also increased the age effect for the four other metrics, bringing the Lighthouse and Cabin errors to uncorrected p = .046 and p = .040 (Table S8).

## Exploratory Analyses

### Directional Bias

Participant mean directions are summarised by age group in Table S13. The clearest directional effect was on LH-Cabin Error, where both groups systematically underestimated the angular separation between the two landmarks by roughly 30° to 40° (younger M = −38.2°, 95% CI [−52.3, −24.2], R = 0.82; older M = −32.5°, 95% CI [−49.3, −15.8], R = 0.86). Confidence intervals also excluded zero for Lighthouse Error in the younger group (M = 13.6°, 95% CI [0.7, 26.4], R = 0.87) and for Start-LH Error in the younger group (M = −10.6°, 95% CI [−18.4, −2.8], R = 0.93). All remaining intervals included zero. Mean resultant lengths ranged from 0.65 to 0.93, indicating that participants within each group were generally consistent in the direction of their bias; because the unit of analysis is the participant mean direction, R indexes agreement across participants rather than the precision of individual pointing responses.

Watson’s U² comparisons of the two groups’ directional distributions were significant before correction for Start Error (U² = 0.193, p = .041) and Lighthouse Error (U² = 0.193, p = .041), but no comparison survived Holm correction across the five metrics (all Holm p ≥ .206). However, because the two groups also differed in concentration, a significant U² is not attributable to a shift in mean direction alone. Block-wise directional summaries are reported in Table S14b.

### Practice Effects (Block 1 vs. 7)

Thirteen participants (6 younger, 7 older) completed a seventh block repeating the path configurations of block 1. The error magnitudes of these participants did not substantially differ from the remainder of the sample in blocks 1-6 (Cohen’s d = −0.26 to 0.23 Table S15a). Error magnitudes decreased numerically on all five metrics between block 1 and block 7 (changes of −9.8° to −23.0°), but no change survived Holm correction (smallest Holm p = .109 for Cabin Error; Table S15).

### Error Coupling

Within-trial coupling between pointing responses was tested at the participant level (Table S16). There was significant directional coupling between the two landmark responses (Lighthouse-Cabin mean r = 0.26, 95% CI [0.16, 0.36], Holm p < .001), as was magnitude coupling for the same pair (mean r = 0.21, 95% CI [0.09, 0.32], Holm p = .002).

In contrast, coupling between the start response and either landmark response was indistinguishable from zero (directional: Start-Lighthouse mean r = −0.01, Holm p = .96; Start-Cabin mean r = −0.02, Holm p = .96; magnitude coupling likewise null). No coupling measure significantly differed between age groups (all Holm ps> .99).

Participants who reported using the sun as a directional cue (n = 15) showed weaker Start-Lighthouse directional coupling than non-users (−0.11 versus 0.04; difference −0.15, 95% CI [−0.27, −0.02], p = .031), although this did not survive Holm correction (Holm p = .093).

### Exploratory Covariates

Sun use, state and trait anxiety, depressive symptoms, and self-reported family history of cognitive impairment were each added one at a time to the primary model (Table S17). No effect survived Benjamini-Hochberg false discovery rate correction across the 25 tests (smallest q = .219). Nominally significant associations, reported here as hypothesis-generating, were smaller Cabin Error magnitude (ratio = 0.72, p = .013) and LH-Cabin Error magnitude (ratio = 0.67, p = .025) among self-reported sun users, and larger Lighthouse Error magnitude (ratio = 1.25, p = .044), Start-LH Error magnitude (ratio = 1.23, p = .036), and LH-Cabin Error magnitude (ratio = 1.45, p = .026) among participants self-reporting a family history of cognitive impairment. The Start Error magnitude age effect was relatively unchanged by every exploratory covariate (ratio = 1.52 to 1.61 against an unadjusted value of 1.54 across observed the group gap).

### Feature ablation

To characterize which of the hand-crafted features carried age-related information, the 28 features were partitioned into six groups by behavioural dimension (angular velocity, 4 features; heading, 4; smoothness and pausing, 4; path length and duration, 6; controller movement, 2; spatial error, 8), and each group was removed in turn. The baseline and all ablated conditions used an untuned support vector classifier (RBF kernel, regularization C and kernel coefficient gamma at their scikit-learn defaults, class weights balanced by group frequency, probabilities Platt-calibrated), evaluated under the same participant-level aggregation and the same leave-one-participant-out scheme throughout, so that only the feature subset varied. The full 28-feature model is the baseline against which each drop is measured (the “None” row of Table S19)

Four of the six groups were informative in the sense that removing them degraded classification (Table S19). The largest single loss came from removing controller movement (−7.8 points), which is notable given that this group comprises only two features. Removing smoothness and pausing (−6.9), heading (−6.3), or angular velocity (−3.0) also reduced BA. Heading and angular velocity were the two informative groups whose individual removal cost the least (−6.3 and −3.0 points, against −7.8 and −6.9 for controller movement and smoothness and pausing), so this pair was tested together to see whether their losses compounded; no other group pairing was tested. Dropping them together did not recover the loss: the resulting 20-feature model reached 65.6% balanced accuracy, matching the larger of the two single-group drops (heading alone) exactly on balanced accuracy, and falling below both single-group drops on AUC (0.694 versus 0.710 and 0.711). The two groups therefore appear to carry overlapping rather than redundant information.

Dropping path length and duration left balanced accuracy essentially unchanged (+0.9 points, within the resolution of a single participant in this sample). Dropping the eight spatial error features produced the largest nominal gain (+1.7 points), accompanied by a considerably larger change in AUC, from 0.733 to 0.818. The AUC change in particular indicates that the error features were actively degrading the ordering of participants by predicted probability rather than merely failing to help. This is consistent with the error-metric findings in Section 3.2, where only Start Error showed a reliable age effect and the four remaining metrics did not, so that six of the eight error features contribute variance without contributing an age-related signal.

This ablation was conducted on the untuned baseline model, which is not among the classifiers reported in the main results; whether the same pattern holds for the tuned model has not been tested on the current N = 54 sample. Adopting a feature subset on the basis of its aggregate performance across all held-out folds would also constitute a selection made with sight of the labels it is subsequently scored against, the same class of leakage corrected in the fusion threshold procedure (Section 2.6). The ablation is therefore reported as a diagnostic of feature-group informativeness for the untuned baseline model and not as a basis for the reported models. No statistical test comparing feature subsets is reported.

## 2. Supplementary Figures

**Supplementary Figure 1.**
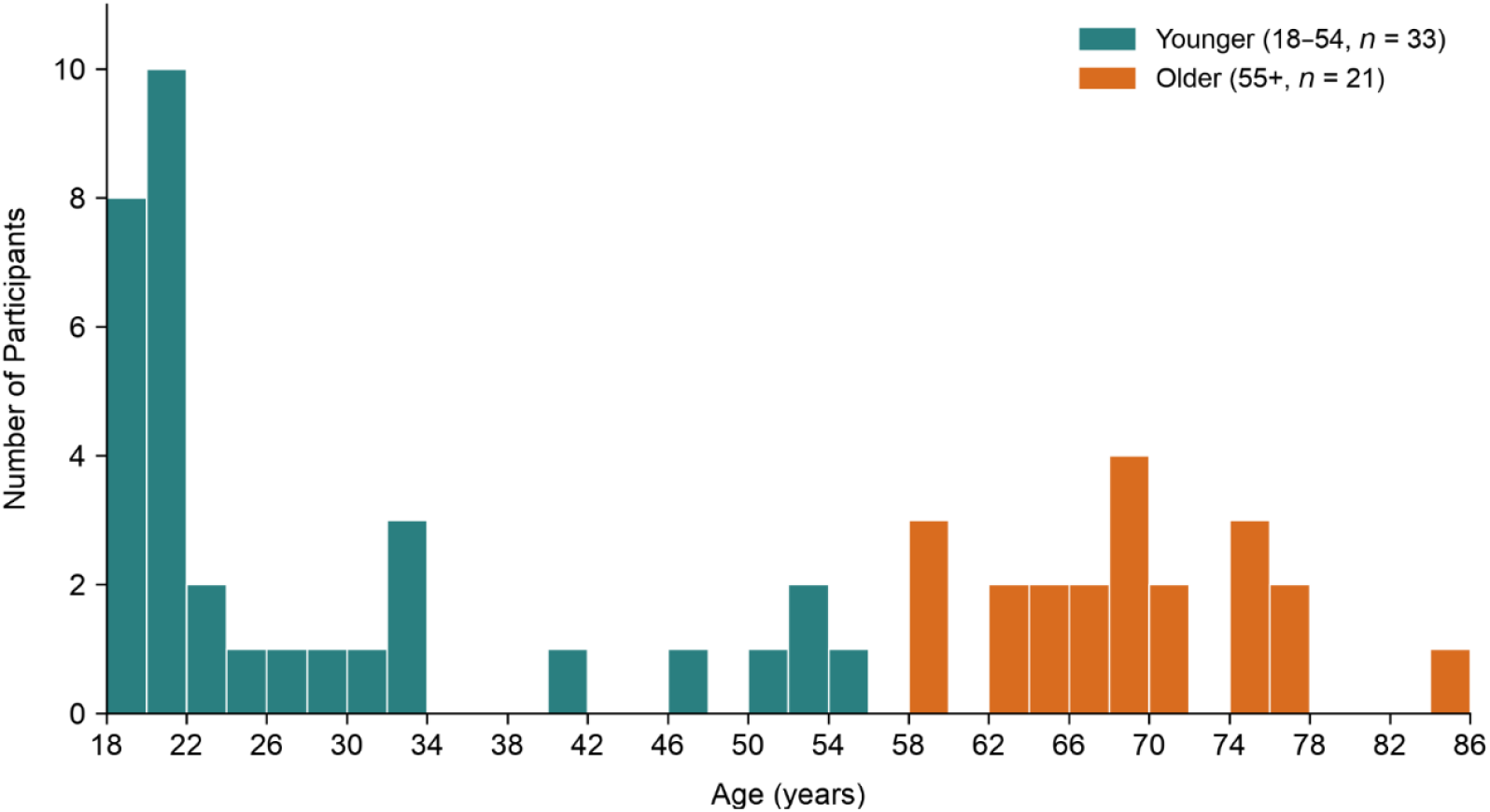
Age Distribution of participants. 54 adults (30 female, 24 male) aged 18 to 84 years (43.30 ± 22.63) were included in the study. Most younger participants were between 18 and 22, having been recruited from the UBC Psychology Department Human Subject Pool. We chose the cutoff between older and younger participants at 55.

**Supplementary Figure 2.**
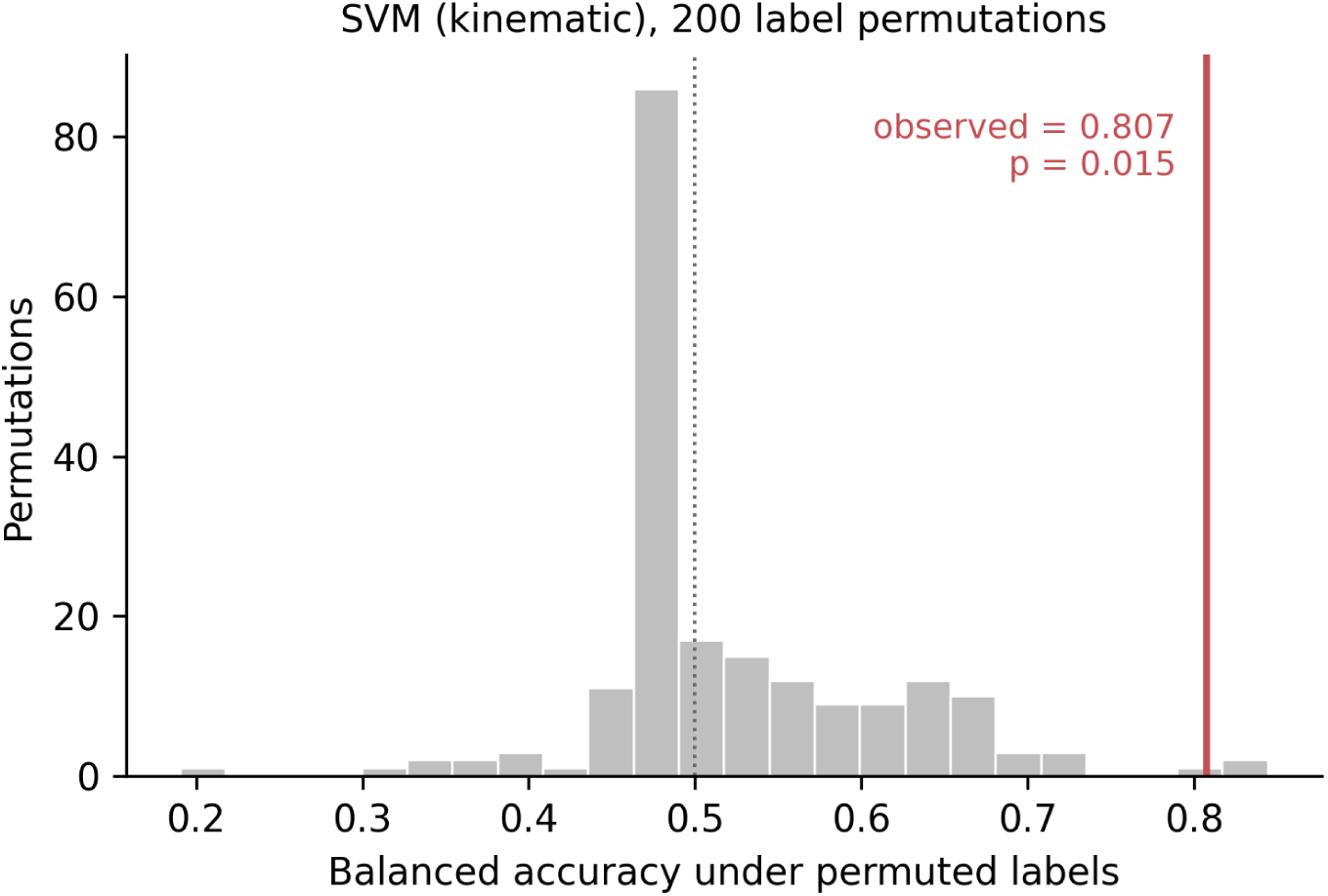
Null distribution of BA for the kinematic SVM under 200 label permutations. Each permutation re-ran the full nested cross-validation including hyperparameter re-selection, and applied the same per-fold Youden threshold rule as the observed analysis. The solid red line marks the observed BA of 80.7%, exceeded only by 2/200 permutations (p=0.015 by the add-one estimator, Phipson and Smyth, 2010). The dotted line marks chance. The feature subset was held fixed across permutations, so this test addresses whether the model exceeds chance, not whether the kinematic subset outperforms the full feature set.

## 3. Supplementary Tables

### Supplementary Table 1 – Attrition Checks

**Table S1.** Blocks 1-3 Performance, Completers vs Non-Completers.

| **Metric** | **Younger, non-completers**<br>**n=1** | **Younger, completers**<br>**n=32** | **Older, non-completers**<br>**n=4** | **Older, completers**<br>**n=17** |
|---|---|---|---|---|
| Start Error | 19.9 | 36.3 (23.6) | 29.1 (10.2) | 59.5 (38.2) |
| LH Error | 52.3 | 56.8 (20.2) | 76.1 (27.4) | 65.1 (21.8) |
| Cabin Error | 56.2 | 62.3 (21.5) | 79.1 (14.5) | 65.4 (22.8) |
| Start-LH Error | 39.5 | 42.1 (13.5) | 62.0 (23.5) | 48.1 (15.3) |
| LH-Cabin Error | 50.0 | 55.6 (32.8) | 55.6 (55.1) | 38.0 (18.7) |

Note. Participant-level mean (SD) angular error magnitude in degrees, in blocks 1-3. There was only one young non-completer, so we report the value. Older non-completers show lower Start Error but higher error on the remaining four metrics than older completers, predicting that the completers-only analysis (Table S6) will inflate the Start Error age effect and attenuate the others.

### Supplementary Table 2 – Model Specification Checks

**Table S2a.** Distribution Family Comparison.

| **Metric** | **n** | **AIC<br>(Gamma)** | **AIC<br>(lognormal)** | **Delta<br>AIC** | **Preferred** |
|---|---|---|---|---|---|
| Start Error | 1,535 | 14573.8 | 14916.1 | -342.3 | gamma |
| LH Error | 1,535 | 15593.7 | 15889.9 | -296.2 | gamma |
| Cabin Error | 1,535 | 15732.5 | 16147.3 | -414.8 | gamma |
| Start-LH Error | 1,535 | 14858.7 | 15148.8 | -290.1 | gamma |
| LH-Cabin Error | 1,535 | 14768.1 | 15028.8 | -260.8 | gamma |

Note. Marginal fits to trial-level absolute error in blocks 1-6. Negative Delta AIC favours the Gamma distribution.

**Table S2b.**
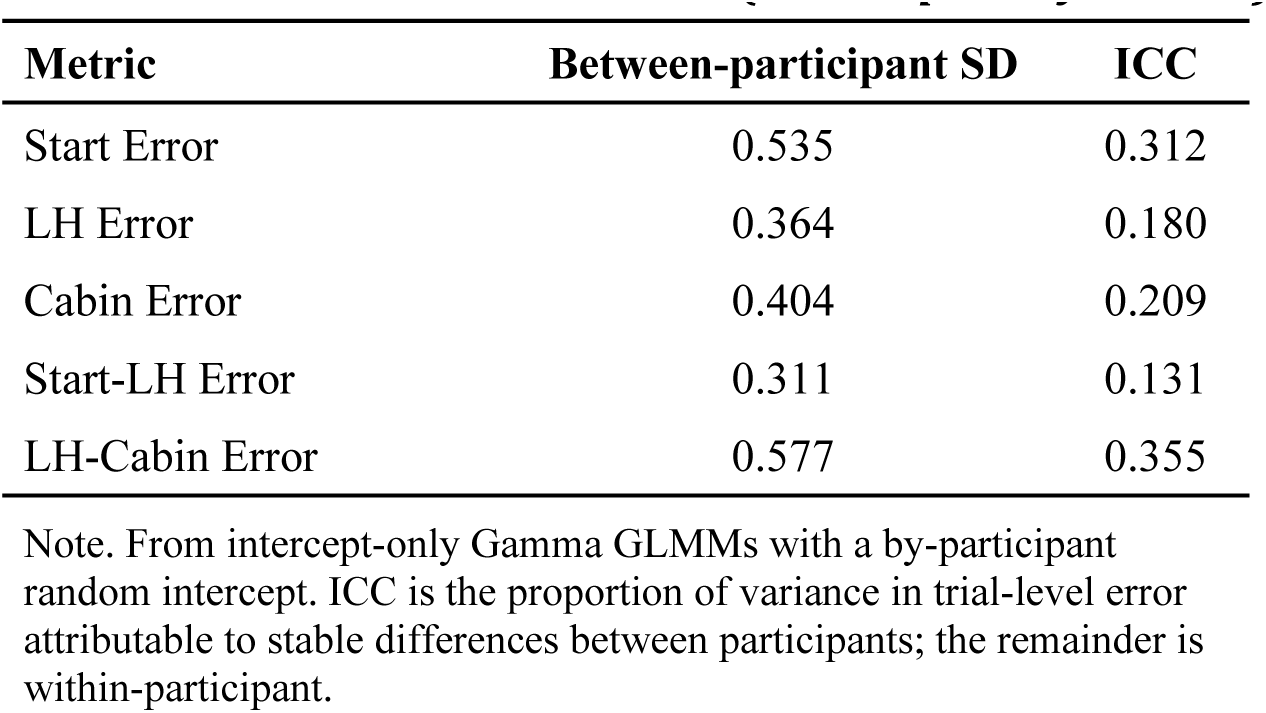
Intraclass Correlation (Intercept-Only Models)

**Table S2c.**
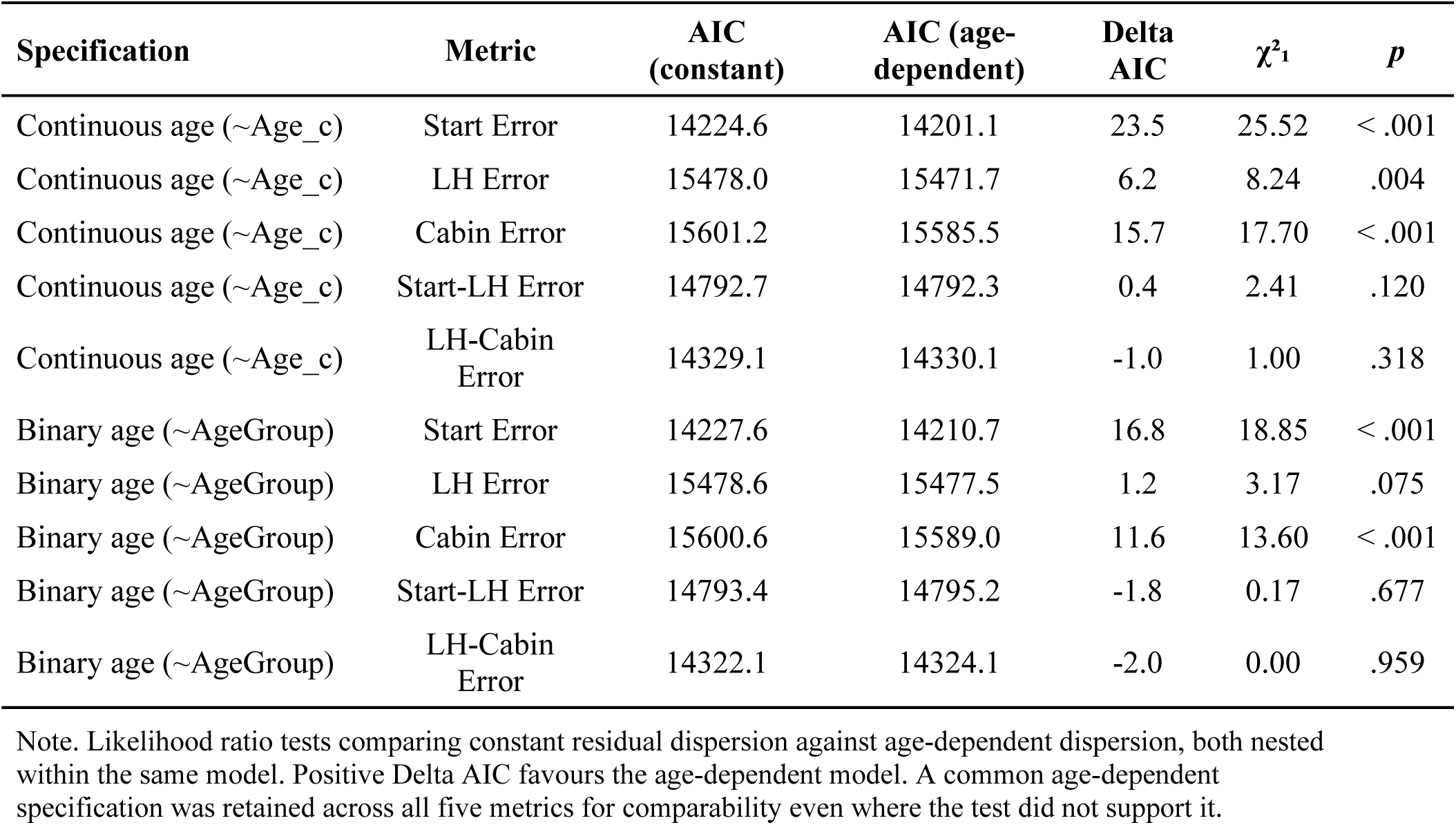
Dispersion Structure Comparison.

| **Specification** | **Metric** | **AIC (constant)** | **AIC (age-dependent)** | **Delta AIC** | **χ<sup>2</sup><sub>1</sub>** | ***p*** |
|---|---|---|---|---|---|---|
| Continuous age (~Age_c) | Start Error | 14224.6 | 14201.1 | 23.5 | 25.52 | < .001 |
| Continuous age (~Age_c) | LH Error | 15478.0 | 15471.7 | 6.2 | 8.24 | .004 |
| Continuous age (~Age_c) | Cabin Error | 15601.2 | 15585.5 | 15.7 | 17.70 | < .001 |
| Continuous age (~Age_c) | Start-LH Error | 14792.7 | 14792.3 | 0.4 | 2.41 | .120 |
| Continuous age (~Age_c) | LH-Cabin Error | 14329.1 | 14330.1 | -1.0 | 1.00 | .318 |
| Binary age (~AgeGroup) | Start Error | 14227.6 | 14210.7 | 16.8 | 18.85 | < .001 |
| Binary age (~AgeGroup) | LH Error | 15478.6 | 15477.5 | 1.2 | 3.17 | .075 |
| Binary age (~AgeGroup) | Cabin Error | 15600.6 | 15589.0 | 11.6 | 13.60 | < .001 |
| Binary age (~AgeGroup) | Start-LH Error | 14793.4 | 14795.2 | -1.8 | 0.17 | .677 |
| Binary age (~AgeGroup) | LH-Cabin Error | 14322.1 | 14324.1 | -2.0 | 0.00 | .959 |

Note. Likelihood ratio tests comparing constant residual dispersion against age-dependent dispersion, both nested within the same model. Positive Delta AIC favours the age-dependent model. A common age-dependent specification was retained across all five metrics for comparability even where the test did not support it.

**Table S2d.** Dispersion Component Coefficients.

| Specification | Metric | Term | *b* | SE | 95% CI | p |
|---|---|---|---|---|---|---|
| Continuous age (~Age_c) | Start Error | Intercept | 0.1493 | 0.0327 | [0.0852, 0.2133] | < .001 |
| Continuous age (~Age_c) | Start Error | Age (per year) | 0.0073 | 0.0014 | [0.0045, 0.0102] | < .001 |
| Continuous age (~Age_c) | LH Error | Intercept | 0.2036 | 0.0328 | [0.1392, 0.2679] | < .001 |
| Continuous age (~Age_c) | LH Error | Age (per year) | 0.0043 | 0.0015 | [0.0014, 0.0072] | .004 |
| Continuous age (~Age_c) | Cabin Error | Intercept | 0.1804 | 0.0328 | [0.1161, 0.2446] | < .001 |
| Continuous age (~Age_c) | Cabin Error | Age (per year) | 0.0063 | 0.0015 | [0.0034, 0.0092] | < .001 |
| Continuous age (~Age_c) | Start-LH Error | Intercept | 0.1049 | 0.0326 | [0.0411, 0.1688] | .001 |
| Continuous age (~Age_c) | Start-LH Error | Age (per year) | 0.0023 | 0.0015 | [-0.0006, 0.0051] | .120 |
| Continuous age (~Age_c) | LH-Cabin Error | Intercept | 0.2045 | 0.0328 | [0.1401, 0.2688] | < .001 |
| Continuous age (~Age_c) | LH-Cabin Error | Age (per year) | -0.0015 | 0.0015 | [-0.0044, 0.0014] | .318 |
| Binary age (~AgeGroup) | Start Error | Intercept | 0.0294 | 0.0408 | [-0.0505, 0.1093] | .471 |
| Binary age (~AgeGroup) | Start Error | Older vs younger | 0.2987 | 0.0679 | [0.1655, 0.4319] | < .001 |
| Binary age (~AgeGroup) | LH Error | Intercept | 0.1523 | 0.0412 | [0.0715, 0.2331] | < .001 |
| Binary age (~AgeGroup) | LH Error | Older vs younger | 0.1217 | 0.0680 | [-0.0116, 0.2551] | .074 |
| Binary age (~AgeGroup) | Cabin Error | Intercept | 0.0797 | 0.0409 | [-0.0005, 0.1600] | .052 |
| Binary age (~AgeGroup) | Cabin Error | Older vs younger | 0.2538 | 0.0681 | [0.1204, 0.3873] | < .001 |
| Binary age (~AgeGroup) | Start-LH Error | Intercept | 0.0914 | 0.0410 | [0.0111, 0.1717] | .026 |
| Binary age (~AgeGroup) | Start-LH Error | Older vs younger | 0.0281 | 0.0673 | [-0.1038, 0.1600] | .676 |
| Binary age (~AgeGroup) | LH-Cabin Error | Intercept | 0.2106 | 0.0414 | [0.1295, 0.2918] | < .001 |
| Binary age (~AgeGroup) | LH-Cabin Error | Older vs younger | -0.0035 | 0.0679 | [-0.1366, 0.1296] | .959 |

Note. Coefficients from the dispersion component of the primary continuous age and binary age models, on the log scale in which they are estimated. This component describes the Gamma dispersion parameter rather than variability in degrees, therefore coefficients are reported untransformed for model transparency and are not interpreted as estimates of response variability. Intercepts give the dispersion parameter at the reference level (mean age for the continuous specification; the younger group for the binary specification). Wald confidence intervals and p-values test whether the dispersion parameter varies with age; they do not index the magnitude or direction of any difference in response variability. Corresponding likelihood ratio tests are reported in Table S2c.

**Table S3.**
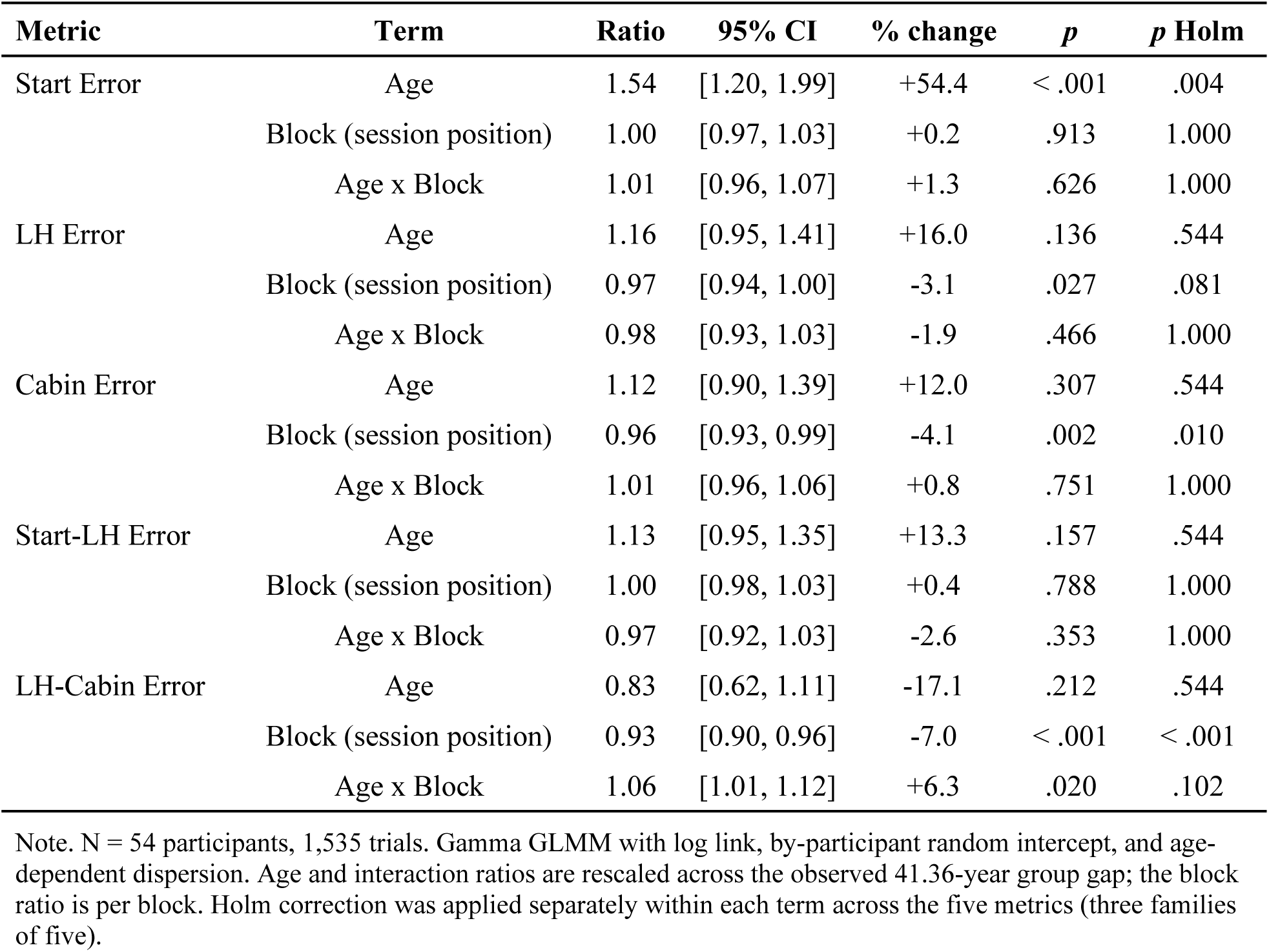
Continuous-Age GLMM: Age, Block, and Interaction Effects.

| **Metric** | **Term** | **Ratio** | **95% CI** | **% change** | ***p*** | ***p* Holm** |
|---|---|---|---|---|---|---|
| Start Error | Age | 1.54 | [1.20, 1.99] | +54.4 | < .001 | .004 |
|  | Block (session position) | 1.00 | [0.97, 1.03] | +0.2 | .913 | 1.000 |
|  | Age x Block | 1.01 | [0.96, 1.07] | +1.3 | .626 | 1.000 |
| LH Error | Age | 1.16 | [0.95, 1.41] | +16.0 | .136 | .544 |
|  | Block (session position) | 0.97 | [0.94, 1.00] | -3.1 | .027 | .081 |
|  | Age x Block | 0.98 | [0.93, 1.03] | -1.9 | .466 | 1.000 |
| Cabin Error | Age | 1.12 | [0.90, 1.39] | +12.0 | .307 | .544 |
|  | Block (session position) | 0.96 | [0.93, 0.99] | -4.1 | .002 | .010 |
|  | Age x Block | 1.01 | [0.96, 1.06] | +0.8 | .751 | 1.000 |
| Start-LH Error | Age | 1.13 | [0.95, 1.35] | +13.3 | .157 | .544 |
|  | Block (session position) | 1.00 | [0.98, 1.03] | +0.4 | .788 | 1.000 |
|  | Age x Block | 0.97 | [0.92, 1.03] | -2.6 | .353 | 1.000 |
| LH-Cabin Error | Age | 0.83 | [0.62, 1.11] | -17.1 | .212 | .544 |
|  | Block (session position) | 0.93 | [0.90, 0.96] | -7.0 | < .001 | < .001 |
|  | Age x Block | 1.06 | [1.01, 1.12] | +6.3 | .020 | .102 |

Note. N = 54 participants, 1,535 trials. Gamma GLMM with log link, by-participant random intercept, and age-dependent dispersion. Age and interaction ratios are rescaled across the observed 41.36-year group gap; the block ratio is per block. Holm correction was applied separately within each term across the five metrics (three families of five).

**Table S4.** Binary-Age GLMM: Age Group, Block, and Interaction Effects.

| Metric | Term | Ratio | 95% CI | % change | *p* | *p* Holm |
|---|---|---|---|---|---|---|
| Start Error | Age group (older vs younger) | 1.52 | [1.14, 2.02] | +51.8 | .004 | .021 |
|  | Block (session position) | 0.99 | [0.96, 1.03] | -0.5 | .785 | 1.000 |
|  | Age group x Block | 1.02 | [0.96, 1.08] | +1.8 | .536 | 1.000 |
| LH Error | Age group (older vs younger) | 1.17 | [0.94, 1.45] | +16.7 | .165 | .661 |
|  | Block (session position) | 0.97 | [0.94, 1.01] | -2.7 | .126 | .377 |
|  | Age group x Block | 0.99 | [0.94, 1.05] | -0.9 | .751 | 1.000 |
| Cabin Error | Age group (older vs younger) | 1.15 | [0.91, 1.47] | +15.4 | .245 | .661 |
|  | Block (session position) | 0.95 | [0.92, 0.99] | -4.9 | .005 | .020 |
|  | Age group x Block | 1.02 | [0.97, 1.08] | +2.2 | .438 | 1.000 |
| Start-LH Error | Age group (older vs younger) | 1.14 | [0.94, 1.39] | +14.4 | .173 | .661 |
|  | Block (session position) | 1.01 | [0.97, 1.05] | +1.1 | .555 | 1.000 |
|  | Age group x Block | 0.98 | [0.92, 1.04] | -1.7 | .572 | 1.000 |
| LH-Cabin Error | Age group (older vs younger) | 0.89 | [0.64, 1.24] | -11.2 | .482 | .661 |
|  | Block (session position) | 0.89 | [0.86, 0.92] | -10.7 | < .001 | < .001 |
|  | Age group x Block | 1.11 | [1.05, 1.18] | +11.2 | < .001 | .001 |

Note. Robustness specification substituting the continuous age classification for binary younger/older (split at 55 years). Ratios are older relative to younger at the session midpoint (block\_c = 0). Holm correction was applied separately within each term across the five metrics. The Age Group x Block interaction for LH-Cabin Error reaches significance here but not under the primary continuous-age specification (Table S3).

**Table S5.** Trial-to-Trial Variability, Completers-Only Panel.

| **Metric** | **Younger SD** | **Older SD** | **Difference [95% CI]** | **Hedges g** | **p** | **p Holm** |
|---|---|---|---|---|---|---|
| Start Error | 28.6 | 37.9 | 9.34 [2.11, 16.30] | 0.77 | .013 | .064 |
| LH Error | 45.7 | 47.3 | 1.60 [-3.94, 7.13] | 0.15 | .613 | 1.000 |
| Cabin Error | 46.4 | 47.0 | 0.54 [-6.41, 6.45] | 0.05 | .881 | 1.000 |
| Start-LH Error | 38.1 | 40.3 | 2.18 [-3.13, 7.55] | 0.22 | .456 | 1.000 |
| LH-Cabin Error | 33.6 | 29.5 | -4.07 [-9.79, 1.60] | -0.42 | .160 | .639 |

Note. Sensitivity analysis repeating Table 4 on the balanced panel of 49 participants (32 younger, 17 older) who completed all 30 analysis trials. Trial-to-trial variability is the standard deviation of absolute error across each participant's trials, in degrees; tabled values are group means of those per-participant standard deviations. Differences are older relative to younger, so a positive difference and a positive Hedges' g indicate greater variability in older adults. P-values are from two-sided Monte Carlo permutation tests (100,000 resamples) randomly re-assigning participants to age groups; confidence intervals are percentile bootstrap (10,000 resamples). p Holm reflects Holm correction across the five metrics. The same panel is used for the completers-only GLMM sensitivity analysis in Table S6.

**Table S6.** Completers-Only Sensitivity Analysis.

| **Specification** | **Metric** | **Full-sample ratio** | **Completers ratio** | **95% CI** | ***p*** | ***p* Holm** | **Direction preserved** |
|---|---|---|---|---|---|---|---|
| Continuous age | Start Error | 1.54 | 1.64 | [1.25, 2.15] | < .001 | .002 | Yes |
|  | LH Error | 1.16 | 1.13 | [0.91, 1.40] | .274 | .821 | Yes |
|  | Cabin Error | 1.12 | 1.09 | [0.86, 1.38] | .493 | .821 | Yes |
|  | Start-LH Error | 1.13 | 1.10 | [0.91, 1.32] | .340 | .821 | Yes |
|  | LH-Cabin Error | 0.83 | 0.81 | [0.59, 1.10] | .178 | .714 | Yes |
| Binary age | Start Error | 1.52 | 1.63 | [1.19, 2.23] | .002 | .011 | Yes |
|  | LH Error | 1.17 | 1.13 | [0.89, 1.43] | .326 | 1.000 | Yes |
|  | Cabin Error | 1.15 | 1.11 | [0.85, 1.46] | .432 | 1.000 | Yes |
|  | Start-LH Error | 1.14 | 1.10 | [0.89, 1.35] | .387 | 1.000 | Yes |
|  | LH-Cabin Error | 0.89 | 0.85 | [0.59, 1.20] | .355 | 1.000 | Yes |

Note. Balanced panel of 49 participants (32 younger, 17 older) contributing all 30 trials in blocks 1-6, against the full sample of 54. Ratios are the age effect only. Continuous-age ratios are rescaled across the full-sample 41.36-year group gap for comparability to the Age Group specification. Holm correction is applied within the five-outcome family of each completers-only model.

### Supplementary Table 7 – Sensitivity: Covariate Adjusted Models

**Table S7a.** Age Effect Before and After Covariate Adjustment.

| **Specification** | **Metric** | **Unadjusted ratio** | **Adjusted ratio** | **95% CI** | ***p*** | ***p* Holm** |
|---|---|---|---|---|---|---|
| Continuous age | Start Error | 1.54 | 1.47 | [1.09, 1.97] | .011 | .055 |
|  | LH Error | 1.16 | 1.17 | [0.93, 1.47] | .188 | .751 |
|  | Cabin Error | 1.12 | 1.17 | [0.91, 1.52] | .221 | .751 |
|  | Start-LH Error | 1.13 | 1.13 | [0.92, 1.38] | .250 | .751 |
|  | LH-Cabin Error | 0.83 | 0.91 | [0.64, 1.29] | .588 | .751 |
| Binary age | Start Error | 1.52 | 1.38 | [1.01, 1.89] | .044 | .221 |
|  | LH Error | 1.17 | 1.15 | [0.90, 1.46] | .267 | .854 |
|  | Cabin Error | 1.15 | 1.19 | [0.91, 1.55] | .213 | .854 |
|  | Start-LH Error | 1.14 | 1.12 | [0.90, 1.38] | .308 | .854 |
|  | LH-Cabin Error | 0.89 | 0.99 | [0.68, 1.43] | .960 | .960 |

Note. Adjustment set: sex (male reference), z-scored years of education, and z-scored Corsi span, entered as main effects. Continuous-age ratios are rescaled across the 41.36-year group gap for comparability to the binary Age Group specification.

**Table S7b.** Covariate Effects in the Adjusted Models.

| **Specification** | **Covariate** | **Metric** | **Ratio** | **95% CI** | ***p*** | ***q* (BH)** |
|---|---|---|---|---|---|---|
| Continuous age | Sex (female) | Start Error | 1.22 | [0.94, 1.60] | .140 | .653 |
|  |  | LH Error | 1.07 | [0.87, 1.32] | .518 | .751 |
|  |  | Cabin Error | 0.99 | [0.79, 1.25] | .959 | .996 |
|  |  | Start-LH Error | 1.07 | [0.89, 1.28] | .499 | .751 |
|  |  | LH-Cabin Error | 1.00 | [0.73, 1.38] | .996 | .996 |
|  | Education (z) | Start Error | 1.02 | [0.87, 1.21] | .777 | .896 |
|  |  | LH Error | 0.93 | [0.82, 1.05] | .253 | .653 |
|  |  | Cabin Error | 0.89 | [0.78, 1.03] | .118 | .653 |
|  |  | Start-LH Error | 0.94 | [0.84, 1.05] | .261 | .653 |
|  |  | LH-Cabin Error | 0.95 | [0.78, 1.15] | .598 | .751 |
|  | Corsi span (z) | Start Error | 0.94 | [0.81, 1.11] | .486 | .751 |
|  |  | LH Error | 0.92 | [0.82, 1.04] | .202 | .653 |
|  |  | Cabin Error | 0.94 | [0.82, 1.07] | .347 | .743 |
|  |  | Start-LH Error | 0.92 | [0.82, 1.03] | .133 | .653 |
|  |  | LH-Cabin Error | 1.05 | [0.87, 1.27] | .601 | .751 |
| Binary age | Sex (female) | Start Error | 1.21 | [0.92, 1.59] | .180 | .653 |
|  |  | LH Error | 1.06 | [0.86, 1.31] | .566 | .653 |
|  |  | Cabin Error | 0.99 | [0.78, 1.25] | .918 | .965 |
|  |  | Start-LH Error | 1.06 | [0.88, 1.27] | .539 | .653 |
|  |  | LH-Cabin Error | 1.01 | [0.73, 1.39] | .965 | .965 |
|  | Education (z) | Start Error | 1.06 | [0.90, 1.25] | .469 | .653 |
|  |  | LH Error | 0.94 | [0.83, 1.06] | .331 | .653 |
|  |  | Cabin Error | 0.90 | [0.79, 1.03] | .139 | .653 |
|  |  | Start-LH Error | 0.95 | [0.85, 1.05] | .319 | .653 |
|  |  | LH-Cabin Error | 0.93 | [0.77, 1.12] | .458 | .653 |
|  | Corsi span (z) | Start Error | 0.93 | [0.79, 1.10] | .393 | .653 |
|  |  | LH Error | 0.92 | [0.81, 1.04] | .172 | .653 |
|  |  | Cabin Error | 0.93 | [0.81, 1.07] | .326 | .653 |
|  |  | Start-LH Error | 0.92 | [0.82, 1.02] | .115 | .653 |
|  |  | LH-Cabin Error | 1.06 | [0.88, 1.29] | .518 | .653 |

Note. Ratios are multiplicative changes per unit of the covariate (one SD for z-scored variables; female relative to male for sex). Benjamini-Hochberg correction was applied across the 15 covariate tests within each age specification.

### Supplementary Table 8 – Sensitivity: Gaming and VR Experience

**Table S8a.** Gaming and VR Experience Sensitivity (Continuous-Age Model)

| **Covariate** | **Metric** | **Age ratio before** | **Age ratio after** | **Age p** | **Covariate ratio** | **95% CI** | **Covariate *p*** | ***q* (BH)** |
|---|---|---|---|---|---|---|---|---|
| Gaming experience (z) | Start Error | 1.47 | 1.26 | .157 | 0.82 | [0.69, 0.98] | .029 | .130 |
|  | LH Error | 1.17 | 1.29 | .046 | 1.13 | [0.99, 1.30] | .078 | .164 |
|  | Cabin Error | 1.17 | 1.34 | .040 | 1.17 | [1.01, 1.36] | .039 | .130 |
|  | Start-LH Error | 1.13 | 1.23 | .073 | 1.11 | [0.98, 1.25] | .095 | .164 |
|  | LH-Cabin Error | 0.91 | 1.11 | .585 | 1.28 | [1.05, 1.57] | .016 | .130 |
| VR experience (z) | Start Error | 1.47 | 1.40 | .032 | 0.92 | [0.78, 1.07] | .279 | .399 |
|  | LH Error | 1.17 | 1.20 | .129 | 1.05 | [0.93, 1.19] | .395 | .494 |
|  | Cabin Error | 1.17 | 1.21 | .169 | 1.05 | [0.91, 1.20] | .494 | .539 |
|  | Start-LH Error | 1.13 | 1.15 | .198 | 1.03 | [0.93, 1.15] | .539 | .539 |
|  | LH-Cabin Error | 0.91 | 0.99 | .960 | 1.17 | [0.97, 1.40] | .098 | .164 |

Note. Gaming and VR experience were each added separately to the adjusted model (sex, education, Corsi span). Age ratio before and after refer to the age effect with and without the interface covariate. Benjamini-Hochberg correction was applied across the pooled 10-test covariate family (2 covariates x 5 metrics) within this age specification. The age p-values are uncorrected. Age ratios are rescaled across the observed 41.36-year group gap.

**Table S8b.** Gaming and VR Experience Sensitivity (Binary-Age Model)

| **Covariate** | **Metric** | **Age ratio before** | **Age ratio after** | **Age p** | **Covariate ratio** | **95% CI** | **Covariate *p*** | ***q* (BH)** |
|---|---|---|---|---|---|---|---|---|
| Gaming experience (z) | Start Error | 1.38 | 1.21 | .234 | 0.81 | [0.68, 0.95] | .010 | .054 |
|  | LH Error | 1.15 | 1.22 | .128 | 1.10 | [0.96, 1.25] | .160 | .286 |
|  | Cabin Error | 1.19 | 1.28 | .074 | 1.14 | [0.99, 1.32] | .075 | .187 |
|  | Start-LH Error | 1.12 | 1.17 | .156 | 1.08 | [0.97, 1.22] | .172 | .286 |
|  | LH-Cabin Error | 0.99 | 1.16 | .443 | 1.29 | [1.06, 1.56] | .011 | .054 |
| VR experience (z) | Start Error | 1.38 | 1.30 | .123 | 0.91 | [0.77, 1.07] | .255 | .364 |
|  | LH Error | 1.15 | 1.19 | .182 | 1.06 | [0.93, 1.20] | .398 | .497 |
|  | Cabin Error | 1.19 | 1.23 | .153 | 1.05 | [0.92, 1.21] | .455 | .505 |
|  | Start-LH Error | 1.12 | 1.15 | .239 | 1.04 | [0.93, 1.16] | .525 | .525 |
|  | LH-Cabin Error | 0.99 | 1.11 | .578 | 1.19 | [0.99, 1.43] | .068 | .187 |

Note. Gaming and VR experience were each added separately to the adjusted model (sex, education, Corsi span). Age ratio before and after refer to the age effect with and without the interface covariate. Benjamini-Hochberg correction was applied across the pooled 10-test covariate family (2 covariates x 5 metrics) within this age specification. The age p-values are uncorrected. Age ratios are older relative to younger at the session midpoint.

### Supplementary Table 9 – Model Diagnostics

**Table S9a.** DHARMa Residual Diagnostics.

| **Specification** | **Metric** | **KS statistic** | **KS *p*** | **Dispersion statistic** | **Dispersion *p*** | **Outlier *p*** | **Quantile *p*** |
|---|---|---|---|---|---|---|---|
| Continuous age | Start Error | 0.053 | < .001 | 0.74 | .006 | 1.000 | .079 |
|  | LH Error | 0.044 | .005 | 0.79 | .004 | .800 | .019 |
|  | Cabin Error | 0.039 | .017 | 0.71 | < .001 | .760 | .002 |
|  | Start-LH Error | 0.037 | .030 | 0.84 | .022 | .780 | .051 |
|  | LH-Cabin Error | 0.060 | < .001 | 0.64 | < .001 | .240 | .243 |
| Binary age | Start Error | 0.052 | < .001 | 0.74 | < .001 | 1.000 | .084 |
|  | LH Error | 0.043 | .007 | 0.78 | .002 | 1.000 | .010 |
|  | Cabin Error | 0.044 | .006 | 0.71 | < .001 | .700 | .004 |
|  | Start-LH Error | 0.039 | .020 | 0.84 | .024 | .500 | < .001 |
|  | LH-Cabin Error | 0.060 | < .001 | 0.64 | < .001 | .180 | .350 |

Note. Scaled residuals computed from 1000 simulations per model. Departures from uniformity were observed for all five metrics under both specifications. These are consistent with the bounded nature of the outcome: absolute angular error cannot exceed 180 degrees, whereas the Gamma distribution has unbounded support. Dispersion statistics below 1.00 indicate that observed variance was smaller than the model predicted.

**Table S9b.** Participant-Aggregated Uniformity Tests.

| **Specification** | **Metric** | **n groups** | **KS *p*** |
|---|---|---|---|
| Continuous age | Start Error | 54 | < .001 |
|  | LH Error | 54 | .003 |
|  | Cabin Error | 54 | < .001 |
|  | Start-LH Error | 54 | .038 |
|  | LH-Cabin Error | 54 | < .001 |
| Binary age | Start Error | 54 | < .001 |
|  | LH Error | 54 | .003 |
|  | Cabin Error | 54 | < .001 |
|  | Start-LH Error | 54 | .022 |
|  | LH-Cabin Error | 54 | < .001 |

Note. Uniformity tests on scaled residuals aggregated within participant, assessing whether any participant is systematically misfit.

**Table S9c.**
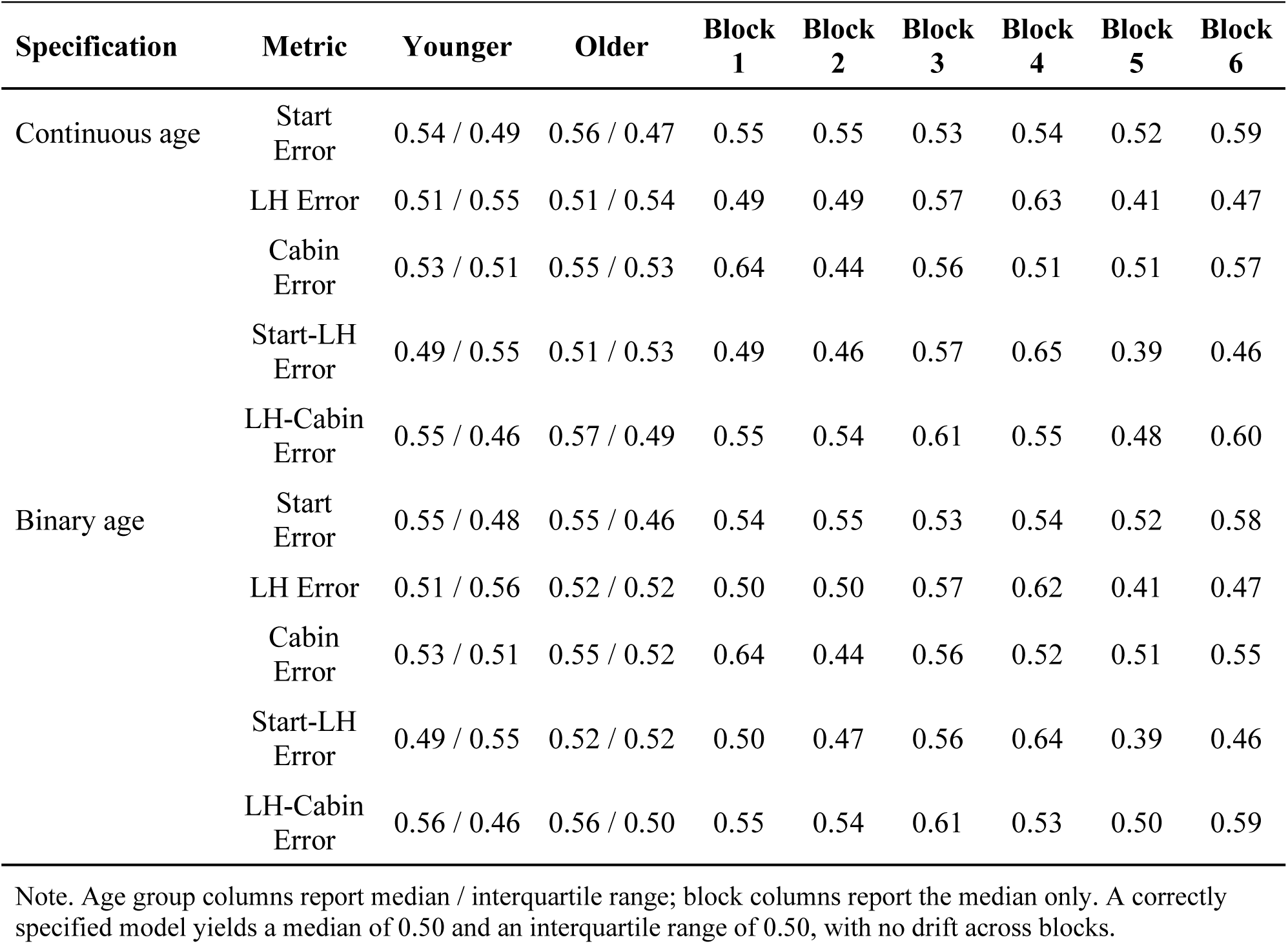
Distribution of Scaled Residuals by Age Group and Block.

| **Specification** | **Metric** | **Younger** | **Older** | **Block 1** | **Block 2** | **Block 3** | **Block 4** | **Block 5** | **Block 6** |
|---|---|---|---|---|---|---|---|---|---|
| Continuous age | Start Error | 0.54 / 0.49 | 0.56 / 0.47 | 0.55 | 0.55 | 0.53 | 0.54 | 0.52 | 0.59 |
|  | LH Error | 0.51 / 0.55 | 0.51 / 0.54 | 0.49 | 0.49 | 0.57 | 0.63 | 0.41 | 0.47 |
|  | Cabin Error | 0.53 / 0.51 | 0.55 / 0.53 | 0.64 | 0.44 | 0.56 | 0.51 | 0.51 | 0.57 |
|  | Start-LH Error | 0.49 / 0.55 | 0.51 / 0.53 | 0.49 | 0.46 | 0.57 | 0.65 | 0.39 | 0.46 |
|  | LH-Cabin Error | 0.55 / 0.46 | 0.57 / 0.49 | 0.55 | 0.54 | 0.61 | 0.55 | 0.48 | 0.60 |
| Binary age | Start Error | 0.55 / 0.48 | 0.55 / 0.46 | 0.54 | 0.55 | 0.53 | 0.54 | 0.52 | 0.58 |
|  | LH Error | 0.51 / 0.56 | 0.52 / 0.52 | 0.50 | 0.50 | 0.57 | 0.62 | 0.41 | 0.47 |
|  | Cabin Error | 0.53 / 0.51 | 0.55 / 0.52 | 0.64 | 0.44 | 0.56 | 0.52 | 0.51 | 0.55 |
|  | Start-LH Error | 0.49 / 0.55 | 0.52 / 0.52 | 0.50 | 0.47 | 0.56 | 0.64 | 0.39 | 0.46 |
|  | LH-Cabin Error | 0.56 / 0.46 | 0.56 / 0.50 | 0.55 | 0.54 | 0.61 | 0.53 | 0.50 | 0.59 |

Note. Age group columns report median / interquartile range; block columns report the median only. A correctly specified model yields a median of 0.50 and an interquartile range of 0.50, with no drift across blocks.

**Table S10.**
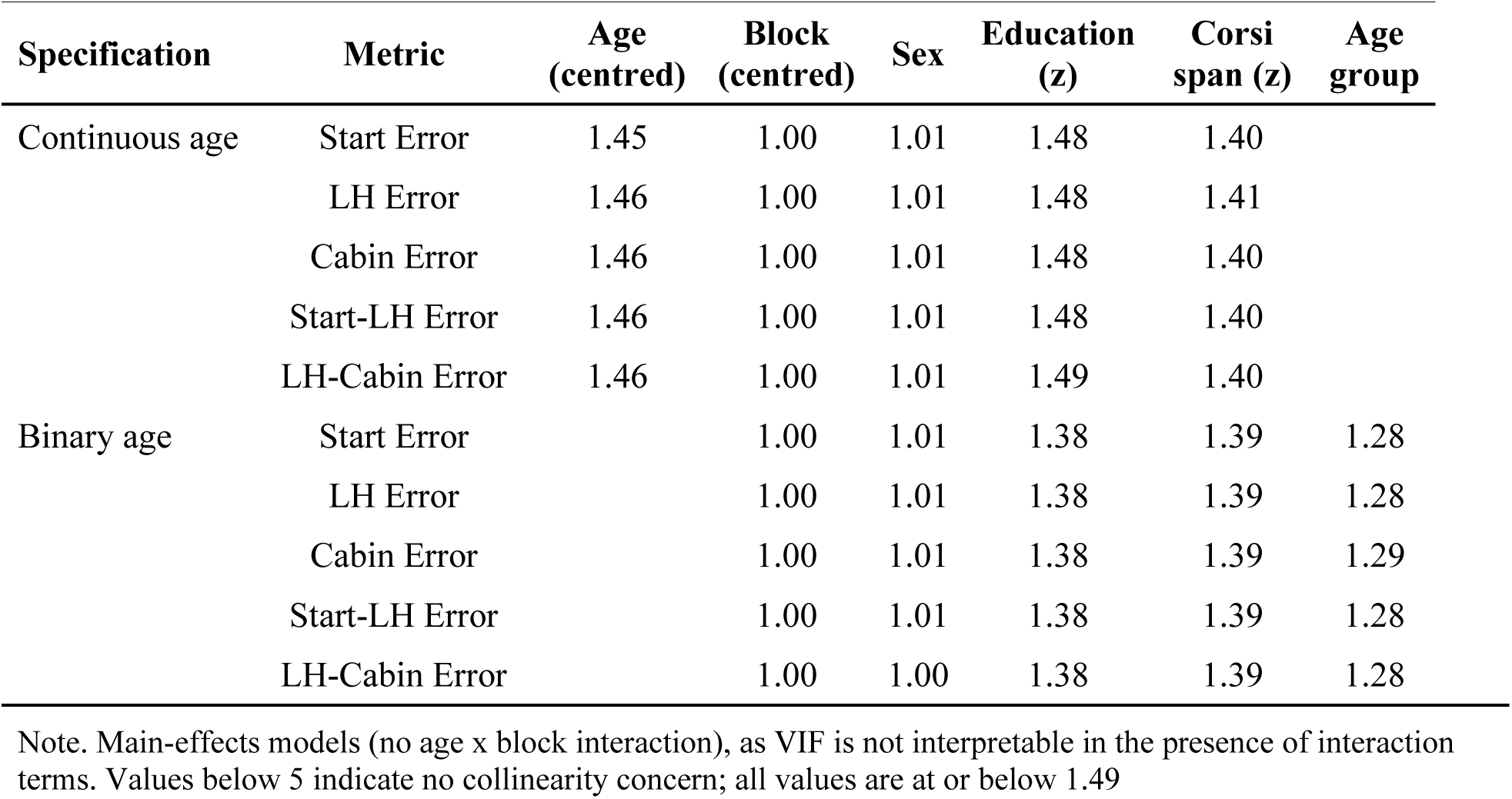
Variance Inflation Factors for the Adjusted Models.

| **Specification** | **Metric** | **Age (centred)** | **Block (centred)** | **Sex** | **Education (z)** | **Corsi span (z)** | **Age group** |
|---|---|---|---|---|---|---|---|
| Continuous age | Start Error | 1.45 | 1.00 | 1.01 | 1.48 | 1.40 |  |
|  | LH Error | 1.46 | 1.00 | 1.01 | 1.48 | 1.41 |  |
|  | Cabin Error | 1.46 | 1.00 | 1.01 | 1.48 | 1.40 |  |
|  | Start-LH Error | 1.46 | 1.00 | 1.01 | 1.48 | 1.40 |  |
|  | LH-Cabin Error | 1.46 | 1.00 | 1.01 | 1.49 | 1.40 |  |
| Binary age | Start Error |  | 1.00 | 1.01 | 1.38 | 1.39 | 1.28 |
|  | LH Error |  | 1.00 | 1.01 | 1.38 | 1.39 | 1.28 |
|  | Cabin Error |  | 1.00 | 1.01 | 1.38 | 1.39 | 1.29 |
|  | Start-LH Error |  | 1.00 | 1.01 | 1.38 | 1.39 | 1.28 |
|  | LH-Cabin Error |  | 1.00 | 1.00 | 1.38 | 1.39 | 1.28 |

Note. Main-effects models (no age x block interaction), as VIF is not interpretable in the presence of interaction terms. Values below 5 indicate no collinearity concern; all values are at or below 1.49

### Supplementary Table 11 – Error Metrics Intercorrelations

**Table S11a.** Spearman Intercorrelations Among Absolute Error Metrics.

| **Metric** | **Start Error** | **LH Error** | **Cabin Error** | **Start-LH Error** | **LH-Cabin Error** |
|---|---|---|---|---|---|
| Start Error | 1.00 |  |  |  |  |
| LH Error | .22 | 1.00 |  |  |  |
| Cabin Error | .17 | .93 | 1.00 |  |  |
| Start-LH Error | .14 | .97 | .90 | 1.00 |  |
| LH-Cabin Error | -.28 | .38 | .50 | .41 | 1.00 |

Note. Participant-level means across blocks 1-6, N = 54. Start-LH Error and Lighthouse Error are near-redundant by construction: both depend on the pointed lighthouse direction, and their absolute values differ only on trials where the pointed lighthouse was placed on the opposite side of the true start direction (or of the direction 180° from it) from the true lighthouse. The correlation pattern suggests approximately three clusters: Start Error; Lighthouse, Cabin and Start-LH Errors; and LH-Cabin Error.

**Table S11b.** Circular-Circular Intercorrelations Among Signed Error Metrics.

| **Metric** | **Start Error** | **LH Error** | **Cabin Error** | **Start-LH Error** | **LH-Cabin Error** |
|---|---|---|---|---|---|
| Start Error | 1.00 |  |  |  |  |
| LH Error | -.25 | 1.00 |  |  |  |
| Cabin Error | .13 | .27 | 1.00 |  |  |
| Start-LH Error | .02 | -.02 | -.01 | 1.00 |  |
| LH-Cabin Error | -.11 | -.08 | -.17 | .86 | 1.00 |

Note. Jammalamadaka-Sarma circular correlations computed on participant mean directions, N = 54.

**Table S12.** Spearman Correlations Among Participant-Level Covariates.

| **Metric** | **Age** | **Education (years)** | **Corsi span** | **PMAT** | **Gaming experience** | **VR experience** | **State anxiety** | **Trait anxiety** | **Depression** |
|---|---|---|---|---|---|---|---|---|---|
| Age | 1.00 |  |  |  |  |  |  |  |  |
| Education (years) | .66 | 1.00 |  |  |  |  |  |  |  |
| Corsi span | -.45 | -.47 | 1.00 |  |  |  |  |  |  |
| PMAT | -.35 | .01 | .06 | 1.00 |  |  |  |  |  |
| Gaming experience | -.50 | -.27 | .29 | .33 | 1.00 |  |  |  |  |
| VR experience | -.48 | -.33 | .38 | .09 | .45 | 1.00 |  |  |  |
| State anxiety | -.38 | -.42 | .25 | -.08 | .18 | .08 | 1.00 |  |  |
| Trait anxiety | -.43 | -.26 | .20 | .11 | .03 | .14 | .38 | 1.00 |  |
| Depression | -.53 | -.40 | .28 | .10 | .26 | .31 | .39 | .69 | 1.00 |

Note. N = 54. Age is correlated with every continuous covariate at \|rho\| >= .35.

**Table S13.** Directional Bias and Age-Group Comparison.

| **Metric** | **n Younger** | **Younger M (R)** | **Younger 95% CI** | **n Older** | **Older M (R)** | **Older 95% CI** | **U<sup>2</sup>** | ***p*** | ***p* Holm** |
|---|---|---|---|---|---|---|---|---|---|
| Start Error | 33 | -5.2 (0.91) | [-14.5, 4.2] | 21 | -19.1 (0.75) | [-39.6, 1.5] | 0.193 | .041 | .206 |
| LH Error | 33 | 13.6 (0.87) | [0.7, 26.4] | 21 | 6.0 (0.65) | [-18.8, 30.8] | 0.193 | .041 | .206 |
| Cabin Error | 33 | 12.7 (0.78) | [-2.6, 27.9] | 21 | 11.5 (0.71) | [-10.8, 33.7] | 0.113 | .211 | .634 |
| Start-LH Error | 33 | -10.6 (0.93) | [-18.4, -2.8] | 21 | -2.6 (0.91) | [-14.4, 9.3] | 0.043 | .809 | 1.000 |
| LH-Cabin Error | 33 | -38.2 (0.82) | [-52.3, -24.2] | 21 | -32.5 (0.86) | [-49.3, -15.8] | 0.035 | .910 | 1.000 |

Note. Values are computed on participant mean directions (N = 54); R indexes the consistency of directional bias across participants rather than the precision of individual pointing responses. Watson's U-squared is a nonparametric omnibus test of whether two circular samples derive from the same distribution; a significant result may reflect a difference in mean direction, in concentration, or in distributional shape. P-values are from 10,000 permutations of the age-group labels, Holm-corrected across the five metrics.

### Supplementary Table 14 – Error Metrics Descriptives by Block

**Table S14a.**
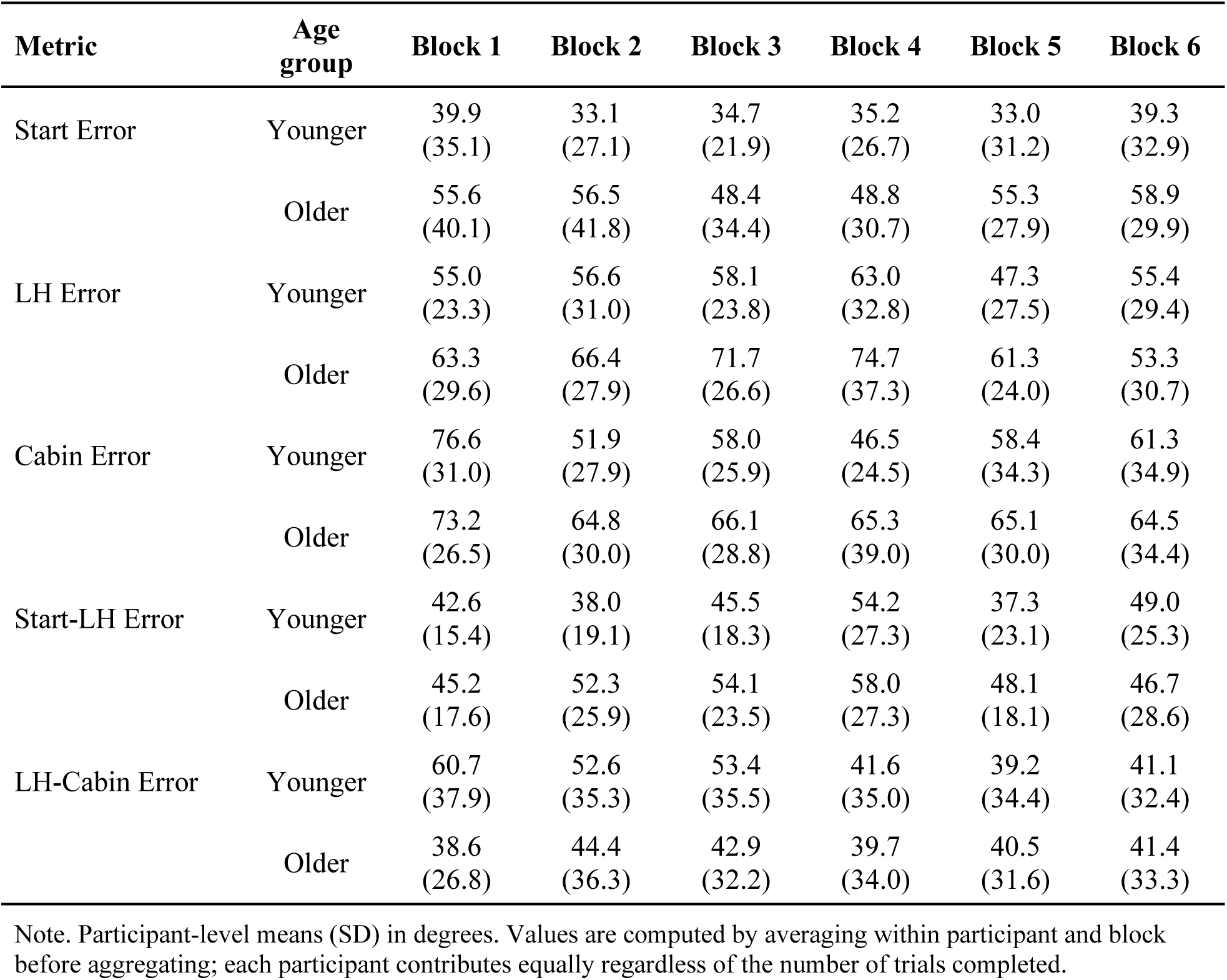
Angular Error Magnitude by Block and Age Group.

**Table S14b.** Directional Bias by Block and Age Group.

| Metric | Age group | Block 1 | Block 2 | Block 3 | Block 4 | Block 5 | Block 6 |
|---|---|---|---|---|---|---|---|
| Start Error | Younger | -15.1 [0.69] | -5.2 [0.86] | 1.8 [0.88] | -4.4 [0.86] | -6.7 [0.84] | -11.0 [0.77] |
|  | Older | -19.0 [0.54] | -9.8 [0.51] | -13.9 [0.65] | 10.5 [0.61] | -15.1 [0.65] | -24.1 [0.67] |
| LH Error | Younger | 28.0 [0.77] | 26.3 [0.69] | 6.2 [0.71] | 36.1 [0.56] | -9.3 [0.74] | -6.5 [0.72] |
|  | Older | -6.2 [0.42] | 23.7 [0.59] | 22.4 [0.52] | -19.3 [0.33] | -21.7 [0.74] | -4.1 [0.70] |
| Cabin Error | Younger | 44.6 [0.36] | 7.4 [0.78] | 35.3 [0.80] | 3.6 [0.81] | 0.9 [0.49] | -0.1 [0.42] |
|  | Older | 16.8 [0.45] | 17.7 [0.56] | 43.4 [0.56] | -15.6 [0.44] | -16.9 [0.52] | -12.5 [0.40] |
| Start-LH Error | Younger | -1.9 [0.89] | -13.6 [0.83] | 0.8 [0.84] | -42.0 [0.78] | -4.4 [0.85] | 5.3 [0.92] |
|  | Older | 22.8 [0.87] | -29.8 [0.74] | -2.8 [0.68] | -38.9 [0.82] | 2.5 [0.79] | 16.9 [0.87] |
| LH-Cabin Error | Younger | -51.9 [0.76] | -45.9 [0.78] | -44.1 [0.79] | -30.9 [0.79] | -28.3 [0.81] | -27.4 [0.81] |
|  | Older | -30.0 [0.89] | -34.0 [0.81] | -35.2 [0.85] | -28.1 [0.81] | -32.5 [0.83] | -31.9 [0.81] |

Note. Mean direction in degrees with mean resultant length (R), computed on participant mean directions within each block. R indexes directional agreement across participants, not within-participant precision.

### Supplementary Table 15 – Exploratory: Block 1 vs 7

**Table S15a.** Blocks 1-6 Performance of Block 7 Recipients.

| **Metric** | **Completed Block 7** | **No Block 7** | **Difference** | **Cohen d** |
|---|---|---|---|---|
| Start Error | 36.8 | 43.7 | -7.0 | -0.26 |
| LH Error | 61.4 | 59.0 | +2.4 | 0.11 |
| Cabin Error | 66.1 | 61.0 | +5.1 | 0.23 |
| Start-LH Error | 48.5 | 46.4 | +2.1 | 0.13 |
| LH-Cabin Error | 50.0 | 44.5 | +5.5 | 0.18 |

Note. Participant-level mean angular error magnitudes in degrees across blocks 1-6, comparing the 13 participants who completed block 7 with the 41 who did not. Block 7 was introduced partway through data collection and offered to all subsequent participants.

**Table S15b.** Within-Participant Change from Block 1 to Block 7.

| **Metric** | **n** | **Block 1 M** | **Block 7 M** | **Change** | **95% CI** | **SD of change** | ***p*** | ***p* Holm** |
|---|---|---|---|---|---|---|---|---|
| Start Error | 13 | 51.8 | 32.0 | -19.8 | [-42.4, 1.1] | 41.6 | .110 | .330 |
| LH Error | 13 | 57.8 | 45.4 | -12.4 | [-28.0, 4.8] | 31.6 | .180 | .360 |
| Cabin Error | 13 | 81.8 | 58.8 | -23.0 | [-39.9, -6.4] | 31.4 | .022 | .109 |
| Start-LH Error | 13 | 46.8 | 34.9 | -12.0 | [-20.7, -2.3] | 17.4 | .033 | .132 |
| LH-Cabin Error | 13 | 51.8 | 42.1 | -9.8 | [-24.3, 5.8] | 28.8 | .240 | .360 |

Note. Exploratory: n = 13 (6 younger, 7 older). Block 7 repeated the path configurations of block 1 at the end of the session, isolating practice from path complexity. Negative change indicates improvement. P-values are from sign-flip permutation tests (all 8192 possible sign assignments); confidence intervals are percentile bootstrap (10,000 resamples). Holm correction is across the five metrics.

### Supplementary Table 16 – Exploratory: Error Coupling

**Table S16a.** Error Coupling Descriptives by Age Group.

| **Pair** | **Age group** | **n** | **Directional M (SD)** | **n positive** | **Magnitude M (SD)** | **n positive** |
|---|---|---|---|---|---|---|
| Start~LH | Younger | 33 | -0.014 (0.224) | 17 | 0.013 (0.206) | 19 |
|  | Older | 21 | 0.008 (0.227) | 11 | -0.001 (0.213) | 11 |
| Start~Cabin | Younger | 33 | -0.040 (0.232) | 12 | 0.023 (0.186) | 19 |
|  | Older | 21 | 0.007 (0.224) | 10 | -0.018 (0.253) | 12 |
| LH~Cabin | Younger | 33 | 0.256 (0.353) | 26 | 0.176 (0.407) | 24 |
|  | Older | 21 | 0.266 (0.430) | 16 | 0.258 (0.462) | 15 |

Note. Coupling was computed within each participant across their trials and then treated as a single observation per participant. Directional coupling is the circular-circular correlation between signed errors; magnitude coupling is the Spearman correlation between absolute errors. N positive is the number of participants with a correlation above zero.

**Table S16b.**
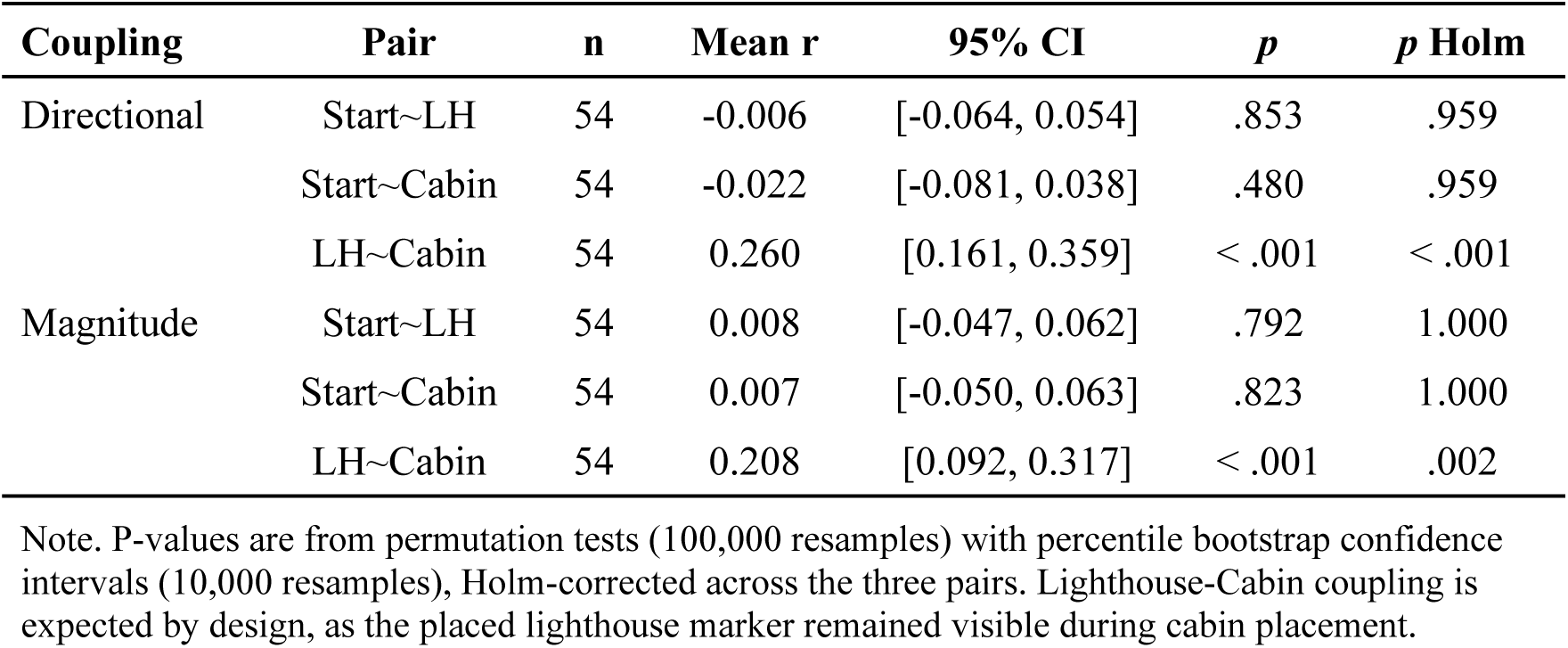
Overall Error Coupling.

**Table S16c.** Error Coupling Group Contrasts.

| **Contrast** | **Coupling** | **Pair** | **Difference** | **95% CI** | ***p*** | ***p* Holm** |
|---|---|---|---|---|---|---|
| Older vs Younger | Directional | Start~LH | 0.022 | [-0.101, 0.142] | .722 | 1.000 |
|  |  | Start~Cabin | 0.047 | [-0.077, 0.170] | .463 | 1.000 |
|  |  | LH~Cabin | 0.009 | [-0.206, 0.222] | .936 | 1.000 |
|  | Magnitude | Start~LH | -0.013 | [-0.126, 0.098] | .819 | 1.000 |
|  |  | Start~Cabin | -0.041 | [-0.163, 0.079] | .491 | 1.000 |
|  |  | LH~Cabin | 0.082 | [-0.157, 0.314] | .495 | 1.000 |
| Sun vs Non-sun | Directional | Start~LH | -0.146 | [-0.273, -0.024] | .031 | .093 |
|  |  | Start~Cabin | -0.071 | [-0.211, 0.061] | .309 | .618 |
|  |  | LH~Cabin | 0.080 | [-0.141, 0.289] | .494 | .618 |
|  | Magnitude | Start~LH | -0.049 | [-0.165, 0.061] | .444 | .444 |
|  |  | Start~Cabin | -0.122 | [-0.250, 0.012] | .063 | .188 |
|  |  | LH~Cabin | 0.199 | [-0.022, 0.405] | .121 | .241 |

Note. Positive differences indicate stronger coupling in the first-named group.

**Table S16d.** Error Coupling Among Self-Reported Sun Users.

| **Coupling** | **Pair** | **n** | **Mean r** | **95% CI** | ***p*** | ***p* Holm** |
|---|---|---|---|---|---|---|
| Directional | Start~LH | 15 | -0.111 | [-0.218, -0.007] | .066 | .131 |
|  | Start~Cabin | 15 | -0.073 | [-0.190, 0.040] | .252 | .252 |
|  | LH~Cabin | 15 | 0.318 | [0.127, 0.485] | .007 | .020 |
| Magnitude | Start~LH | 15 | -0.028 | [-0.122, 0.055] | .596 | .596 |
|  | Start~Cabin | 15 | -0.082 | [-0.195, 0.034] | .197 | .394 |
|  | LH~Cabin | 15 | 0.351 | [0.177, 0.502] | .003 | .009 |

Note. Coupling was computed within each participant across their trials and then treated as a single observation per participant. Directional coupling is the circular-circular correlation between signed errors; magnitude coupling is the Spearman correlation between absolute errors. Group contrasts were suppressed where any cell held fewer than 10 participants. P-values are from permutation tests (100,000 resamples) with percentile bootstrap confidence intervals (10,000 resamples), Holm-corrected across the three pairs. N = 15. Age contrasts are suppressed, as both groups had fewer than 10 sun users.

### Supplementary Table 17 – Exploratory: Covariates

**Table S17a.** Exploratory Covariate Effects (Continuous-Age Model)

| **Covariate** | **Metric** | **Ratio** | **95% CI** | ***p*** | ***q*** **(BH)** | **Age ratio in model** | **Age ratio unadjusted** |
|---|---|---|---|---|---|---|---|
| Sun use | Start Error | 1.07 | [0.78, 1.47] | .663 | .776 | 1.52 | 1.54 |
|  | LH Error | 0.84 | [0.66, 1.07] | .154 | .427 | 1.20 | 1.16 |
|  | Cabin Error | 0.72 | [0.56, 0.93] | .013 | .219 | 1.20 | 1.12 |
|  | Start-LH Error | 0.85 | [0.69, 1.05] | .142 | .427 | 1.17 | 1.13 |
|  | LH-Cabin Error | 0.67 | [0.47, 0.95] | .025 | .219 | 0.90 | 0.83 |
| State anxiety (z) | Start Error | 1.02 | [0.88, 1.19] | .760 | .826 | 1.57 | 1.54 |
|  | LH Error | 1.06 | [0.95, 1.19] | .294 | .603 | 1.21 | 1.16 |
|  | Cabin Error | 1.08 | [0.95, 1.23] | .239 | .572 | 1.17 | 1.12 |
|  | Start-LH Error | 1.06 | [0.96, 1.18] | .252 | .572 | 1.18 | 1.13 |
|  | LH-Cabin Error | 1.06 | [0.89, 1.26] | .505 | .664 | 0.86 | 0.83 |
| Trait anxiety (z) | Start Error | 1.03 | [0.89, 1.20] | .662 | .776 | 1.58 | 1.54 |
|  | LH Error | 0.99 | [0.88, 1.11] | .834 | .869 | 1.15 | 1.16 |
|  | Cabin Error | 0.96 | [0.84, 1.09] | .494 | .664 | 1.08 | 1.12 |
|  | Start-LH Error | 0.98 | [0.88, 1.09] | .683 | .776 | 1.12 | 1.13 |
|  | LH-Cabin Error | 0.94 | [0.79, 1.11] | .455 | .664 | 0.79 | 0.83 |
| Depression (z) | Start Error | 1.01 | [0.86, 1.18] | .940 | .940 | 1.55 | 1.54 |
|  | LH Error | 1.05 | [0.93, 1.19] | .394 | .657 | 1.22 | 1.16 |
|  | Cabin Error | 1.06 | [0.92, 1.21] | .431 | .664 | 1.18 | 1.12 |
|  | Start-LH Error | 1.05 | [0.95, 1.17] | .337 | .603 | 1.19 | 1.13 |
|  | LH-Cabin Error | 1.17 | [0.98, 1.40] | .087 | .312 | 0.95 | 0.83 |
| Family history | Start Error | 0.86 | [0.65, 1.15] | .317 | .603 | 1.61 | 1.54 |
|  | LH Error | 1.25 | [1.01, 1.55] | .044 | .221 | 1.09 | 1.16 |
|  | Cabin Error | 1.27 | [0.99, 1.62] | .056 | .232 | 1.05 | 1.12 |
|  | Start-LH Error | 1.23 | [1.01, 1.49] | .036 | .221 | 1.07 | 1.13 |
|  | LH-Cabin Error | 1.45 | [1.04, 2.00] | .026 | .219 | 0.75 | 0.83 |

Note. Each covariate was added one at a time to the primary model. Ratios are multiplicative changes per unit of the covariate (one SD for z-scored variables; TRUE relative to FALSE for binary variables). Benjamini-Hochberg correction was applied across all 25 tests within this age specification. The final two columns show the age effect in each model against its unadjusted value. These analyses are exploratory, and no effect survived correction. Age ratios are rescaled across the observed 41.36-year group gap.

**Table S17b.** Exploratory Covariate Effects (Binary-Age Model)

| **Covariate** | **Metric** | **Ratio** | **95% CI** | ***p*** | ***q***<br>**(BH)** | **Age ratio**<br>**in model** | **Age ratio**<br>**unadjusted** |
|---|---|---|---|---|---|---|---|
| Sun use | Start Error | 1.12 | [0.82, 1.54] | .484 | .649 | 1.49 | 1.52 |
|  | LH Error | 0.85 | [0.67, 1.08] | .193 | .536 | 1.20 | 1.17 |
|  | Cabin Error | 0.73 | [0.57, 0.94] | .015 | .201 | 1.22 | 1.15 |
|  | Start-LH Error | 0.86 | [0.70, 1.07] | .176 | .536 | 1.17 | 1.14 |
|  | LH-Cabin Error | 0.65 | [0.46, 0.92] | .016 | .201 | 0.95 | 0.89 |
| State anxiety (z) | Start Error | 0.99 | [0.85, 1.15] | .906 | .943 | 1.51 | 1.52 |
|  | LH Error | 1.05 | [0.94, 1.18] | .395 | .649 | 1.20 | 1.17 |
|  | Cabin Error | 1.07 | [0.95, 1.21] | .277 | .649 | 1.20 | 1.15 |
|  | Start-LH Error | 1.05 | [0.95, 1.16] | .335 | .649 | 1.17 | 1.14 |
|  | LH-Cabin Error | 1.08 | [0.91, 1.28] | .367 | .649 | 0.93 | 0.89 |
| Trait anxiety (z) | Start Error | 1.00 | [0.86, 1.17] | .951 | .951 | 1.52 | 1.52 |
|  | LH Error | 0.98 | [0.87, 1.10] | .736 | .836 | 1.15 | 1.17 |
|  | Cabin Error | 0.96 | [0.84, 1.09] | .494 | .649 | 1.12 | 1.15 |
|  | Start-LH Error | 0.97 | [0.88, 1.08] | .611 | .754 | 1.12 | 1.14 |
|  | LH-Cabin Error | 0.96 | [0.81, 1.14] | .634 | .754 | 0.86 | 0.89 |
| Depression (z) | Start Error | 0.99 | [0.84, 1.16] | .877 | .943 | 1.50 | 1.52 |
|  | LH Error | 1.05 | [0.93, 1.19] | .427 | .649 | 1.23 | 1.17 |
|  | Cabin Error | 1.06 | [0.93, 1.22] | .381 | .649 | 1.23 | 1.15 |
|  | Start-LH Error | 1.05 | [0.95, 1.17] | .348 | .649 | 1.21 | 1.14 |
|  | LH-Cabin Error | 1.20 | [1.00, 1.44] | .046 | .205 | 1.07 | 0.89 |
| Family history | Start Error | 0.89 | [0.67, 1.20] | .456 | .649 | 1.56 | 1.52 |
|  | LH Error | 1.25 | [1.01, 1.55] | .041 | .205 | 1.10 | 1.17 |
|  | Cabin Error | 1.26 | [0.99, 1.60] | .061 | .217 | 1.08 | 1.15 |
|  | Start-LH Error | 1.23 | [1.01, 1.49] | .036 | .205 | 1.08 | 1.14 |
|  | LH-Cabin Error | 1.39 | [1.00, 1.93] | .049 | .205 | 0.81 | 0.89 |

Note. Each covariate was added one at a time to the primary model. Ratios are multiplicative changes per unit of the covariate (one SD for z-scored variables; TRUE relative to FALSE for binary variables). Benjamini-Hochberg correction was applied across all 25 tests within this age specification. The final two columns show the age effect in each model against its unadjusted value. These analyses are exploratory, and no effect survived correction. Age ratios are older relative to younger at the session midpoint (block\_c = 0).

**Table S18.** Start Error Magnitude Age Effect Across Model Specifications.

| Specification | n participants | n trials | Ratio | 95% CI | p | p Holm |
|---|---|---|---|---|---|---|
| Unadjusted (primary) | 54 | 1,535 | 1.54 | [1.20, 1.99] | < .001 | .004 |
| Completers only | 49 | 1,444 | 1.64 | [1.25, 2.15] | < .001 | — |
| Adjusted (sex, education, Corsi) | 54 | 1,535 | 1.47 | [1.09, 1.97] | .011 | — |
| Adjusted + gaming experience | 54 | 1,535 | 1.26 | [0.92, 1.72] | .157 | — |
| Adjusted + VR experience | 54 | 1,535 | 1.40 | [1.03, 1.90] | .032 | — |

Note. All rows report the Age_c coefficient rescaled across the observed 41.36-year group gap. The p column is uncorrected: rows 2-5 are robustness checks on the single primary hypothesis tested in row 1. The p Holm column reports the confirmatory correction from Table 3 and applies only to the primary unadjusted row. Gaming and VR experience are fitted as separate models, each adjusted for sex, education, and Corsi span; their own covariate effects are BH-corrected in Table S8a. The completers-only estimate is larger than the full-sample estimate, consistent with the attrition analysis showing that withdrawn older adults had lower Start Error before withdrawal; the full-sample estimate is retained as primary. Results for the other four metrics appear in Tables S6-S8.

**Table S19.** Drop-One Feature Group Ablation (Untuned SVM)

| **Condition Dropped** | **n features** | **BA** | **Sensitivity** | **Specificity** | **AUC** | **Δ BA** |
|---|---|---|---|---|---|---|
| Controller movement | 26 | 64.1% | 0.524 | 0.758 | 0.680 | −7.8 |
| Smoothness and pausing | 24 | 64.9% | 0.571 | 0.727 | 0.691 | −6.9 |
| Heading | 24 | 65.6% | 0.524 | 0.788 | 0.710 | −6.3 |
| Heading + Angular velocity | 20 | 65.6% | 0.524 | 0.788 | 0.694 | -6.3 |
| Angular Velocity | 24 | 68.8% | 0.619 | 0.758 | 0.711 | -3.0 |
| None (Full feature set) | 28 | 71.9% | 0.619 | 0.818 | 0.733 | 0.0 |
| Path length and duration | 22 | 72.7% | 0.667 | 0.788 | 0.723 | +0.9 |
| Spatial error | 20 | 73.6% | 0.714 | 0.758 | 0.818 | +1.7 |

N = 54, trial cutoff 16, untuned SVM (RBF kernel, balanced class weights), leave-one-participant-out cross-validation. ΔBA is the change in BA in percentage points relative to the full 28-feature set. Rows are ordered by ΔBA.

